# Focus on the breath: Brain decoding reveals internal states of attention during meditation

**DOI:** 10.1101/461590

**Authors:** H.Y. Weng, J.A. Lewis-Peacock, F.M. Hecht, M.R. Uncapher, D.A. Ziegler, N.A.S. Farb, V. Goldman, S. Skinner, L.G. Duncan, M.T. Chao, A. Gazzaley

**Author notes:** Correspondence should be addressed to H.Y.W. 1545 Divisadero St, Suite 300 San Francisco, CA 94115.

## Abstract

Meditation practices are used to cultivate internally-oriented attention to bodily sensations, which may improve health via cognitive and emotion regulation of bodily signals. However, it remains unclear how meditation impacts internal attention states due to lack of measurement tools that can objectively assess mental states during meditation practice itself, and produce time estimates of internal focus at individual or group levels. To address these measurement gaps, we tested the feasibility of applying multi-voxel pattern analysis (MVPA) to single-subject fMRI data to (1) learn and recognize internal attentional (IA) states relevant for meditation during a directed IA task, and (2) decode or estimate the presence of those IA states during an independent meditation session. Within a mixed sample of experienced meditators and novice controls (N=16), we first used MVPA to develop single-subject brain classifiers for 5 modes of attention during an IA task in which subjects were specifically instructed to engage in one of five states (i.e., meditation-related states: breath attention, mind wandering, and self-referential processing, and control states: attention to feet and sounds). Using standard cross-validation procedures, MVPA classifiers were trained in five of six IA blocks for each subject, and predictive accuracy was tested on the independent sixth block (iterated until all block volumes were tested, N=2160). Across participants, all five IA states were significantly recognized well above chance (>41% vs. 20% chance). At the individual level, IA states were recognized in most participants (87.5%), suggesting that recognition of IA neural patterns may be generalizable for most participants, particularly experienced meditators. Next, for those who showed accurate IA neural patterns, the originally trained classifiers were then applied to a separate meditation run (10-min) to make an inference about the percentage time engaged in each IA state (breath attention, mind wandering, or self-referential processing). Preliminary group-level analyses demonstrated that during meditation practice, participants spent more time attending to breath compared to mind wandering or self-referential processing. This paradigm established the feasibility of using MVPA classifiers to objectively assess mental states during meditation at the participant level, which holds promise for improved measurement of internal attention states cultivated by meditation.

---

Mind-body practices such as meditation are increasingly practiced by the public to improve health (Clarke and Stussman 2018), and train internal qualities of attention to the body that include sustained focus, nonjudgment, and compassion (Kabat-Zinn 2008; Gunaratana 2010; Lutz et al. 2015). Through practicing these attentional qualities to bodily sensations such as the breath, meditation practices may strengthen interoception (awareness of internal bodily sensations; Farb *et al.*, 2015; Khalsa *et al.*, 2017), cognitive processes (including sustained attention, cognitive monitoring, and meta-awareness; Lutz *et al.*, 2008; Dahl *et al.*, 2015; Tang *et al.*, 2015), and emotion regulation (less judgment and more equanimity with internal experiences; Chambers *et al.*, 2009; Desbordes *et al.*, 2015). With practice, these skills may lead to better monitoring and regulation of physical, emotional, and social processes, contributing to improved health decision-making and behaviors (Farb *et al.* 2015; Khalsa *et al.* 2017). However, interoceptive processes trained through meditation cannot be directly observed during practice because they are internal and fluctuate among various mental states (such as attention or inattention to the body; Van Dam *et al.*, 2018). While previous studies assess neural activation during meditation to identify networks present at the aggregate level, currently no measure uses neural data to objectively assess whether attention is indeed focused on the body or not during meditation practice itself.

In other modalities of mental training, such as working memory training with external visual stimuli, the trained skills can be *directly observed and measured* in real-time with quantifiable metrics such as working memory performance (Klingberg 2010). In addition, subsequent *transfer effects* to other cognitive skills can be assessed (that are related but not directly trained), such as improvements in working memory in another sensory modality or inhibition (Klingberg 2010). Thus, relationships between skills gained directly from training can be associated with transfer effects. Currently, interventions that train internally-oriented attention (such as meditation and yoga) have been studied mostly through transfer effects (i.e. downstream effects on external attention, emotion regulation, or well-being; Van Dam *et al.*, 2018), largely because we lack measures that can objectively assess the focus of internal attention during practice itself. The field therefore does not currently have a parallel measure to working memory performance, or metric of interoceptive focus or stability, that is both objective and unobtrusive to practice and could provide metrics such as proportion time attending (or not) to the breath during meditation practice. With these metrics, we could more precisely understand how cultivating qualities of internal attention transfers to other psychological processes and more global states such as mental and physical health.

A measure that could directly assess the meditative process would be able to track various mental states as they fluctuate through time. For example, in a core practice of focused attention to the breath, attention is focused on sensations of the breath, until distracted by other internal or external stimuli, and then nonjudgmentally returned to the breath. Even in this simple practice, distinct mental states may occur and dynamically fluctuate over time: the object of attention (breath or distractions), level of meta-awareness (awareness of object of attention), as well as attitudinal qualities such as nonjudgment, kindness, and curiosity (Hasenkamp *et al.* 2012; Lutz *et al.* 2015). Previous studies have mapped out neural networks associated with components of this process using fMRI, identifying greater activation in networks involved in interoception (Farb, Segal, and Anderson 2013; Fox *et al.* 2016) and executive functioning (Brefczynski-Lewis *et al.*, 2007; Fox *et al.*, 2016), and decreased activation of the Default Mode Network (Brewer *et al.*, 2011; Fox *et al.*, 2016), which is engaged during mind wandering and self-referential processing (Andrews-Hanna, Smallwood, and Spreng 2014; Christoff et al. 2016).

Traditional univariate fMRI analysis focuses on mean regional changes in brain activity and therefore usually requires the collapsing of data across many time points to improve signal estimation. The downside of this approach, particularly for the study of meditation, is that such data averaging obscures the fluctuating nature of distinct mental states, such as interoception, mind wandering, and self-referential processing. These measurement issues can be addressed by instead applying multivariate multi-voxel pattern analysis (MVPA; Norman *et al.*, 2006; Haxby, 2012), which uses pattern recognition technology to (i) distinguish and recognize neural patterns associated with external or internal attention, and (ii) then apply these learned brain patterns to decode or estimate the presence of various mental states in a separate task. In this way, MVPA uses objective brain data to “read the mind” during tasks where the mental states are otherwise inaccessible (Norman *et al.* 2006; Haxby 2012). For example, MVPA of fMRI data has been used to reliably differentiate the attentional status of two items held in working memory across an 8-sec memory delay (Lewis-Peacock and Postle 2012). On a trial-by-trial basis, these discrete neural measurements of internal attention have been linked to the precision of behavioral responses on short-term recognition tests (Emrich et al. 2013) and to recognition confidence in tests of long-term memory (Lewis-Peacock and Norman 2014).

In this proof-of-principle study, we aimed to produce quantifiable metrics of interoceptive attention by integrating MVPA methodology to study internal attention states relevant for meditation. We developed the EMBODY framework (**E** valuating **M** ultivariate Maps of **Body** Awareness), where MVPA is applied to neural data to (i) learn and recognize internal attention (IA) states important for breath-focused meditation, and (ii) decode or estimate the presence of those IA states during an independent meditation period. These decoded IA states could then be used to estimate the percentage time engaged in attention or inattention to the breath. Notably, MVPA is often applied using a within-subjects approach, with more data collected from each person so that MVPA classifiers can learn and decode brain patterns that are participant-specific (Norman et al. 2006; Haxby 2012). Similar to what is commonly done in the visual sciences, this approach concentrates experimental power and high-powered tests of effects at the individual level (Smith and Little 2018), and the generalizability of the findings can be assessed by examining the proportion of the subjects in which MVPA is reliable.

We tested the feasibility of the EMBODY framework within 16 participants in 3 steps (**Fig. 1**). In **Step 1**, we tested whether IA brain patterns (breath, mind wandering, self-referential processing) could be reliably learned and distinguished above chance levels by MVPA classifiers. In a directed internal attention task, participants were specifically instructed to engage in one of five states (i.e., meditation-related states: breath attention, mind wandering, and self-referential processing, and control states: attention to feet and sounds), and MVPA was applied to develop single-subject brain classifiers for five modes of attention. Using standard cross-validation procedures, MVPA classifiers were trained in five of six IA blocks for each subject, and predictive accuracy was tested on the independent sixth block (iterated until all block volumes were tested, N=2160). In **Step 2**, for participants who showed accurate IA neural patterns, we applied the originally trained classifiers to a separate meditation run (10-min) to decode or classify IA brain states in a separate 10-min period of breath-focused meditation (N=600). In **Step 3**, we used these classified IA states to make an inference about attention states during meditation, including percentage time engaged in breath-focus, mind wandering, or self-referential processing. Finally, we conducted preliminary assessments of construct validity by associating these brain-derived attention metrics with 1) subjective reports of internal attention within each subject, and 2) attention profiles across the group during the breath-focused meditation period, where we hypothesized participants would spend more time attending to the breath vs. other mental states.

**Figure 1.**
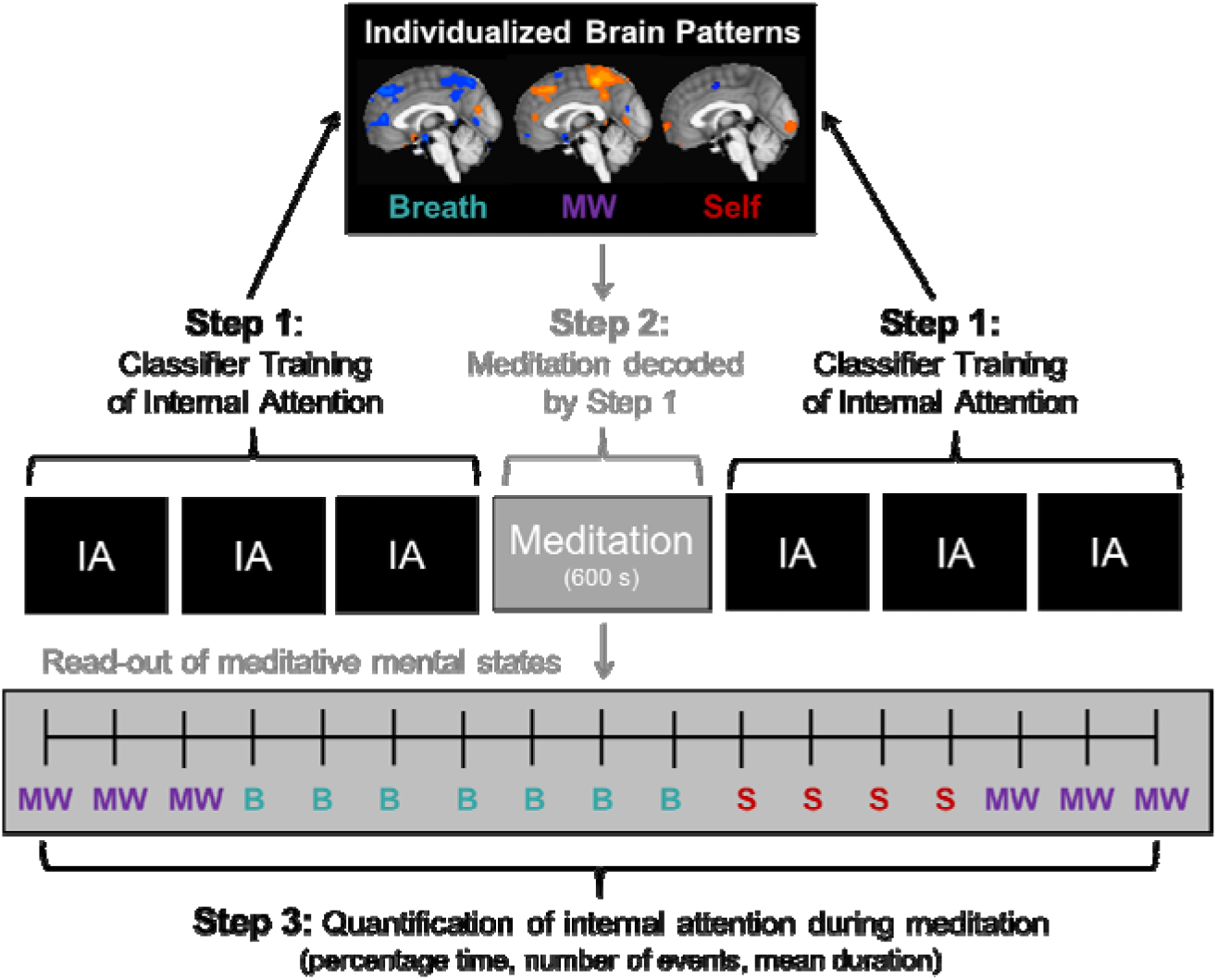
**EMBODY Framework: E**valuating **M**ultivariate Maps of **Body** Awareness to measure internal attention states during meditation. **Step 1. Brain pattern classifier training**. Machine learning algorithms are trained in fMRI neural patterns associated with internal mental states in the Internal Attention (IA) task. IA is directed via auditory instructions to pay attention with eyes closed to the breath, mind wandering (MW), self-referential processing (Self), and control conditions of attention to the feet and ambient sounds (see **Fig. 2**). Individualized brain patterns for each participant are learned using n-1 cross-validation with 6 blocks of the IA task (volume N=2160). **Step 2. Meditation period classification**. Neural patterns are collected during a 10-min meditation period (in this case, focused attention to the breath; administered in the middle of 6 IA blocks), and are decoded by multi-voxel pattern analysis (MVPA) using the unique brain patterns learned in Step 1. Meditation is decoded second-by-second into mental states of attention to breath (B), mind wandering (MW), or self-referential processing (S), producing an estimate of distinct and fluctuating mental states during meditation. **Step 3. Quantification of internal attention during meditation.** From the temporal read-out of meditative mental states in Step 2, attention metrics during meditation can be quantified and estimated including percentage time spent in each mental state, number of times engaged in each mental state (“events”), and mean duration spent in each mental state.

## METHOD

### General framework and approach

We tested the feasibility of the EMBODY framework, which uses MVPA applied to fMRI data to learn and decode mental states during meditation, producing novel individual-level metrics of internal attention during meditation. At a participant-specific level, we first tested whether MVPA classifiers could distinguish between brain patterns of internal attention relevant for meditation significantly above chance levels (**Step 1**), by directing participants to change their focus of internal attention using the Internal Attention (IA) task (**Fig. 2a**). To maximize this within-subjects approach, we collected more data within each participant (2160 brain patterns for classifier training and testing, 432/condition). We tested this framework in 8 meditators and 8 controls (an adequate N for within-subjects MVPA studies; Norman *et al.*, 2006) and included individuals from both groups because a) meditators are more likely to produce distinct brain patterns from consistent practice in directing and sustaining internal attention, and b) novices are the population most studied in clinical intervention studies. To further test whether MVPA classifier accuracy of internal attention was meaningful, we associated accuracy with within-subject subjective ratings of attention.

**Figure 2.**
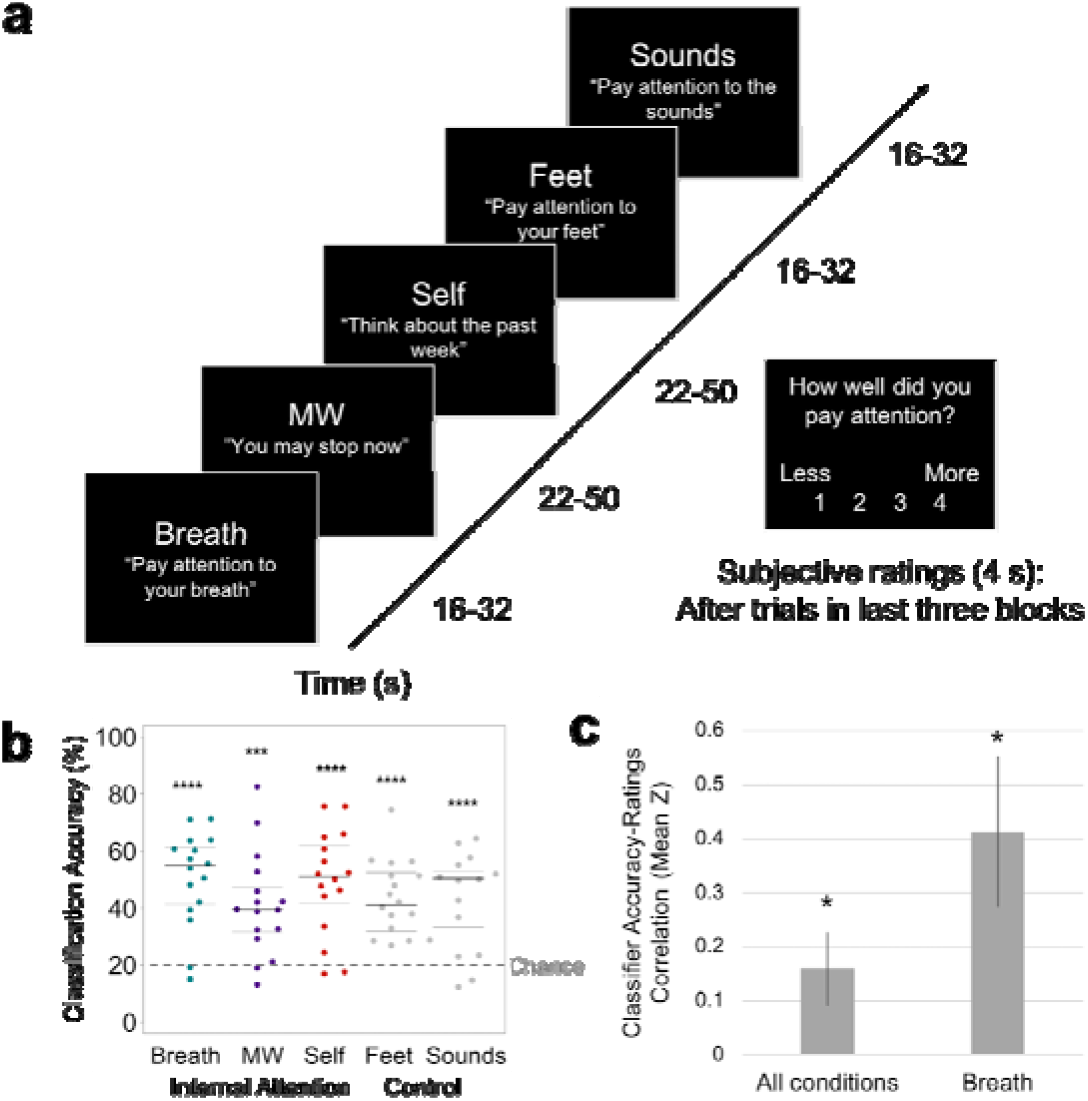
EMBODY Step 1: Classifier training of internal mental states. **(a)** Internal Attention (IA) task. With eyes closed, participants were directed via 2-s auditory instructions to pay attention to five internal mental states for brief time periods (16-50s). The IA task directed attention to three mental states relevant for breath meditation (Breath, MW, and Self), and to two control mental states (attention to the Feet [another area of the body] and ambient MRI Sounds [consistent external distractor]). Example auditory instructions are displayed in quotes. MW was induced by instructing participants to stop paying attention and let their minds attend to whatever they wanted. Conditions were randomized over six IA blocks in four orders, with 72s of data collected from each condition in each block (total 432s/condition). For the last half of IA task trials, subjective ratings of attention were collected after each trial (except MW) using a button box (1=less, 4=more). **(b)** From the IA task, the prediction accuracy of the classifier for identifying internal states of attending to the Breath, MW, and Self, and control conditions of attending to the Feet and Sounds. Beeswarm plots present each data point, the median (bold black line), and ±25^th^ percentile range (gray lines) of the mean prediction accuracy for all data in each condition (N=432) across all subjects. Statistical significance was determined by a one-sample two-sided *t*-test against theoretical chance-level for classification of 5 categories (20%, denoted by dashed line). *** *t*_15_=4.65, *p*<0.001, **** *t*s_15_>5.67, *p*s<0.0001. **(c)** Mean *z*-scores representing the within-subject correlation between trial-level classifier training accuracy and subjective ratings of attention (administered during the last half of IA task trials) for all conditions (except MW) and breath trials only. Error bars indicate standard error of the mean. * *p*<0.05.

By first establishing that MVPA classifiers could reliably distinguish and identify internal attention brain states in Step 1, we could then apply these learned brain patterns to objectively decode the continuous focus of attention using the neural data from a separate 10-min meditation period (600 novel brain patterns, **Step 2**). This is a common application of MVPA, where classifiers learn neural patterns from distinct categories in one task (and classifier accuracy is validated using within-task cross-validation), which are then used to estimate information in a separate task (across-task decoding where classifier training from the first task is used to inform classifier decisions in the second task; Norman *et al.*, 2006), and the EMBODY framework extends this approach into decoding IA states during meditation. We then further analyzed the classifier decisions in **Step 3**, where we computed novel metrics of attention during meditation, including estimating the percentage time attending to the breath or engaging in mind wandering or self-referential processing. This thus served our main measurement goal of *estimating the interoceptive focus or stability* during meditation practice for each individual. Finally, to assess construct validity and inform future research, we preliminarily characterized the meditation metrics at the group level, and tested whether participants attended longer to breath vs. other mental states during meditation.

### Participants

Participants were medically and psychiatrically healthy adults age 25-65, non-smokers, and MRI-compatible. Meditators were recruited from Bay Area meditation centers through flyers, online postings, and word of mouth. They reported consistent meditation practice in the past 5 years (≥90 min/week, ≥14 days of silent retreat practice, at least half of practice on breath and bodily sensations). Control participants were recruited through flyers and online postings, had not engaged in regular meditation or other mind-body practices (>20 min at least twice weekly), and were age (within 5 years) and gender-matched to meditators.

Participants included 8 meditators (1 female, 1 non-binary person, 6 male, mean age = 38.4 [range 28-61], race/ethnicity: 6 White, 2 multiracial) and 8 matched novice control participants (mean age = 38.3 [range 25-63], race/ethnicity: 6 White, 1 Asian, 1 Latinx/Hispanic; see **Table S1** in **Supplemental Information [SI]** for full demographics). Average lifetime meditation practice in meditators was 3495 hours (range 509-6590; **Table S2**). Two additional novices were excluded for inability to align images due to excessive movement and incorrect gender-matching to a meditator.

### Procedure

Eligibility was assessed by online questionnaire and phone interview. Participants were consented, trained in MRI task procedures, and then completed a 2-hour MRI protocol. They were paid $65 for participation and ≤$20 for travel. All participants provided written informed consent approved by the Institutional Review Board of the University of California, San Francisco. The study was registered at clinicaltrials.gov (NCT03344081).

### fMRI Paradigm

#### Overall Framework

The EMBODY Framework used multi-voxel pattern analysis (MVPA; Norman *et al.*, 2006) with fMRI data to decode the focus of internal attention during meditation in 3 steps: 1) participant-specific brain patterns were trained for internal attention states relevant for breath meditation (n=2160), 2) brain patterns learned from Step 1 were applied to a 10-min period of breath meditation to estimate the focus of internal attention for each data point (600s), and 3) metrics of attention during meditation were computed from the decoded brain states (**Fig. 1**).

#### Step 1 data: Internal Attention (IA) task

fMRI data from the IA task were used to train a machine learning classifier to learn neural patterns associated with five internal mental states. To create training data that closely resembled brain activity during meditation, participants’ eyes remained closed, and the only stimulus change was their internal focus of attention. Neural patterns associated with breath, mind wandering, and self-referential processing were chosen to be most relevant for decoding the meditation period, which modeled the intended focus of meditation (breath) and two common distractors (mind wandering and self-referential processing). Neural patterns associated with attention to feet (another body area) and awareness of ambient sounds (consistent MRI sounds) were chosen as control conditions to improve classification specificity of the desired brain states.

Participants received randomized 2-s auditory instructions to pay attention to 1) sensations of the breath (Breath), 2) sensations of the feet (Feet), 3) to stop paying attention and let their minds go wherever they would like (mind wandering or MW), 4) self-referential processing regarding the past, present, and future (Self), and 5) ambient sounds (Sounds; **Fig. 2a**). For breath-focus, attention was maintained where they felt the breath most strongly (e.g., nose, throat, chest). For self-referential processing, participants generated 5 events from the past week, and 5 events that would occur in the next week during the training session. Instructions were also presented visually, which participants could briefly view as a reminder. Six blocks were administered from one of four randomized stimulus order sets.

Each block contained 20s of baseline (black screen) at the beginning and end, and consisted of 13 trials/block, resulting in 72s/condition within each block (balanced across 5 conditions). This yielded 2160 training brain patterns over the entire experiment, with 432 patterns/condition. Trial durations ranged from 16-32s for attending to breath, feet, and sounds (3 rials/block each; every even-numbered trial length was randomized and administered twice/condition across the experiment), and 22-50s in MW and Self (allowing more time for these states to occur, 2 trials/block each, covering most of the duration range across the experiment). In the last three IA blocks, participants subjectively rated how well they paid attention after each trial using a button box (*How well did you pay attention?* 1=less attention, 4=more attention), and were encouraged to use the full range of responses.

#### Step 2 data: Meditation Period

Participants engaged in 10 min of breath-focused meditation in 2 blocks, administered between the 6 IA blocks. The meditation period was split into two blocks (4 and 6 min) to help control participants stay engaged in the task. Participants were instructed to pay attention to sensations of the breath, and if their minds wandered, to return attention to the breath. For each block, they received a 6-s instruction at the beginning, and a 2-s reminder to pay attention 60s before the end. After the meditation period, participants verbally rated the percentage time they paid attention to the breath and thoughts for each block.

### Data acquisition

Experiments were run using E-Prime (Psychology Software Tools). Neuroimaging data were acquired with a 3T MRI scanner (Siemens Prisma) using a 64-channel head coil. A high-resolution 1×1×1 mm MPRAGE T1 anatomical scan was acquired for offline spatial registration. Functional images were acquired using a multiband gradient-echo EPI sequence (2.4×2.4×2.4 mm, TE/TR = 30.2 ms/1 s, FOV=220 mm, 92×92 matrix, 56 slices, multiband acceleration=4, TR=1.0s; Auerbach *et al.*, 2013) that covered most of the brain.

### EMBODY fMRI data analyses: machine learning

#### fMRI preprocessing

Data were preprocessed in AFNI (Cox 1996), and were slice time corrected, aligned and motion-corrected to the first volume of the first EPI scan, and linearly de-trended in native space, respectively using 3dTshift, 3dAllineate, 3dvolreg, 3dDetrend. See **SI** and **Tables S3-S4** for control analyses using head motion and respiration data.

#### Step 1 machine learning: Distinguishing neural patterns of internal attention (within-task cross-validation)

Pattern classification analyses were conducted using MVPA (Norman *et al.*, 2006; Princeton MVPA Toolbox https://github.com/PrincetonUniversity/princeton-mvpa-toolbox), in conjunction with in-house software using Matlab (MathWorks) and Python (for post-processing of meditation period classifications in Steps 2-3). Using wholebrain preprocessed fMRI signal in native space, a pattern classifier was trained separately for each participant for trial periods from each condition (Breath, MW, Self, Feet, and Sounds; TR=1.0s, 432s/condition) using penalized logistic regression with L2 regularization and a penalty parameter of 0.01 (which prevents over-fitting by punishing large weights during classifier training; Duda *et al.*, 2000). Condition labels were shifted in time by 6s to account for hemodynamic lag. A binary logistic regression (1 vs. the others) was run for each of the 5 conditions, resulting in continuous classifier evidence values (0-1) for each condition at each time point in the experiment (**Fig. S1**). The condition that was assigned the highest evidence value yielded the categorical decision from the classifier (see **SI** and **Table S9** for alternate analyses of classifier evidence and decisions). We evaluated classification accuracy by performing k-fold cross-validation analysis, training on five blocks of data (fMRI task runs) and testing on the novel sixth block. The training blocks were then rotated, and a new block of data was tested until all six blocks of data had been classified (2160 decisions).

Classification accuracy for each condition was computed for each participant (the percentage out of 432 accurate decisions output by the machine learning classifier). Group-level accuracy for each condition was tested with a one-sample *t*-test vs. 20% (theoretical chance level for 5 conditions), and the effect size was estimated with Cohen’s *D*. Individual-level accuracy was tested with a Chi-square test determining whether the number of accurate vs. inaccurate decisions in each condition were significantly above chance distribution (87 vs. 345). Individuals that showed above-chance accuracy in 2/3 categories for Breath, MW, and Self conditions were used for subsequent analyses including decoding meditation states (all 8 meditators and 6/8 controls).

#### IA ratings and classifier accuracy

Attention ratings were collected for the second half of trials (33/39 trials, excluding MW trials where no rating was administered). To estimate the correlation of classifier accuracy and ratings within each subject, a Pearson’s correlation was computed between trial-level classifier accuracy and subjective ratings of attention. To test the strength of correlations across the group, each individual *r*-value was transformed using a Fisher *r*-to-*Z* transform, and the group mean *Z*-score was tested vs. 0 using a one-sample *t*-test. Because this task was designed to measure breath attention, we also examined accuracy-rating correlations specifically in Breath trials (n=9).

#### Individualized brain pattern importance maps

Classifier importance maps were computed for each participant using classifier weight information which identifies which voxels were most important in distinguishing neural patterns of Breath, MW, and Self (McDuff, Frankel, and Norman 2009). To identify voxels that distinguished between conditions which were not due to differences in head motion, the analyses were conducted with fMRI data where motion variables were covaried out. We identified voxels with “positive importance” (both the weight and *z*-scored average activation value are positive) and voxels with “negative importance” (both the weight and *z*-scored average activation value are negative). Note that this approach identifies voxels which aid classifier distinction between mental states, and does not test for differences in average activity like standard univariate analyses (Haxby 2012). For display purposes, each individual’s importance maps were non-linearly warped to the MNI152 2mm template using FSL (FNIRT; Smith *et al.*, 2004), smoothed with an 8mm Gaussian kernel, converted to *z*-scores (across voxels), and thresholded at ±2 SD to identify the most important voxels for each condition. See **SI** for group-level importance map analyses.

#### Step 2 machine learning: Decoding the internal focus of attention during breath meditation (across-task decoding)

Individualized brain patterns learned from Step 1 were applied to the 10-min meditation period to decode the internal focus of attention. The meditation period consisted of a completely independent dataset, and these classifiers were not influenced in any way by the previous Step 1 cross-validation analyses. The classifier was trained with all five mental states from the IA task (2160 total brain patterns) and decoded with the three states that were most relevant for breath-focused meditation: Breath, MW, and Self. For each data point during meditation (n=600, excluding instruction periods), the classifier output a categorical decision of whether internal focus was on the Breath, MW, or Self (as well as continuous evidence values for each mental state). This produced a continuous estimate of mental states during meditation over time.

To ignore spurious measurements of brain states that may fluctuate from one time point to the next, we focused our analyses on relatively stable periods. We defined a “mental event” as the classification of three or more consecutive time points for a given category. To facilitate this, we smoothed the data such that a single incongruous decision between two events of the same type (e.g., MW event–Self decision–MW event) were relabeled according to the category of the surrounding events (e.g., Self=>MW; average data points smoothed=1.3%, SD=0.41). Events were then quantified as 3 or more consecutive decisions of the same category, excluding any data that did not meet these criteria (average data excluded = 15.7%, SD = 4.82). See **SI** for additional analyses with no smoothing function and varying mental event lengths (2 and 4s).

#### Step 3: Quantify internal attention metrics during meditation

From the mental state classifications from Step 2, novel metrics of internal attention during meditation were computed for each participant. For each mental state, *percentage time engaged, number of events, mean duration of events*, and *variability* (SD) of event duration were computed. Data were preliminarily analyzed at the group level by testing for differences in metrics between conditions (Breath, MW, Self) with a one-way ANOVA. To test our main hypotheses that breath-focused meditation would result in differences between Breath vs. other mental states, significant results were followed up with planned pair-wise *t*-tests of Breath vs. MW and Breath vs. Self. Data were analyzed in SPSS (v. 24), figures were created with R, and brain maps were displayed using AFNI or FSLview.

## RESULTS

### Step 1: Distinguishing neural patterns of internal attention

The first aim of the EMBODY framework was to test whether MVPA applied to within-subject fMRI data could recognize individualized neural patterns associated with internal attention states important for breath meditation (**Fig. 2a**). Across all participants, each attentional state yielded a distinct neural signature (all classification accuracies>41% vs. 20% chance for 5 categories, *p*s<0.001; **Fig. 2b**). Furthermore, each attentional state was distinguished at more than twice chance levels, including the brain patterns most relevant for breath meditation (breath=50.5%, mind wandering=41.2%, self-referential processing=49.0%; *t*s_15_>4.65, *p*s<0.001, Cohen’s *d*s>1.16) and the control conditions (feet=43.2%, sounds=43.7%, *t*s_15_>5.67, *p*s<0.0001, Cohen’s *d*s>1.41; **Fig. 2b**; see **Table S5** for classifier confusion matrix.

The breath meditation-relevant brain patterns were reliably classified in 14/16 or 87.5% participants (at least 2/3 *p*s<0.001 for breath, mind wandering, and self-referential processing, **Table S7).** This included all 8 meditators and 6 of 8 novices, all of whom were used in subsequent analyses. Within-subject correlations of trial-level classification accuracy and subjective ratings of internal attention (across all conditions except mind wandering) showed a positive association overall across the group (mean *Z*=0.16, one-sample *t*_13_=2.35, *p*=0.035; **Fig. 2c**). Because the task was designed to specifically measure breath attention during meditation, we also examined associations between accuracy and ratings in breath trials only and found a higher mean correlation (mean *Z* =0.41, *t*_13_=2.96, *p*=0.011; **Fig. 2c**).

#### Distributed brain patterns contributing to accurate IA classification

Classifier importance maps identified the voxels most important in distinguishing between the attentional states (McDuff, Frankel, and Norman 2009), which were distributed throughout the brain and unique for each participant (**Fig. 3a**). For initial characterization of brain regions that supported classification and were common across individuals, a frequency map was computed representing the sum of individual importance maps (**Fig. 3b**). This indicated that no brain region was important for all 14 participants in any mental state (maximum frequency≤10, **Fig. 3b**, **Fig. S2a-b**), and frequency histograms showed that most voxels were important for only 1-3 participants (**Fig. S2c-e**). See **Table S7** for preliminary identification of brain regions where voxels were important in a higher frequency of participants (N≥5).

**Figure 3.**
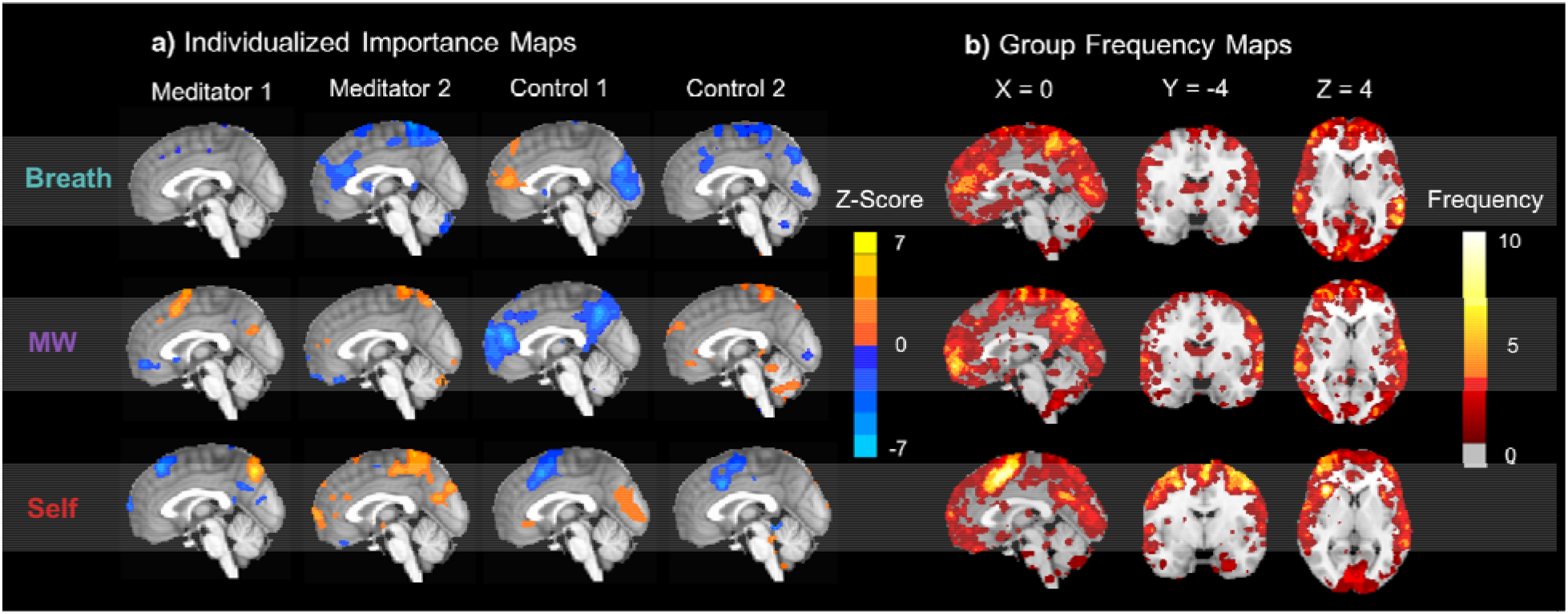
Classifier importance maps representing voxels that accurately distinguish internal mental states. **a)** Subject-level importance maps showing individualized brain patterns representing voxels that are important for distinguishing neural signatures of attention to the Breath, MW, and Self (X=0). For each task condition, importance values were computed by multiplying each voxel’s classifier weight for predicting the condition and the average activation during the condition (McDuff, Frankel, and Norman 2009). The maps were thresholded at ±2 SD and displayed on the MNI152 template to identify the most important voxels for each participant. Orange importance voxel indicate positive *z*-scored average activation values, and blue importance voxels indicate negative *z*-scored average activation values. **b)** For initial characterization of brain regions that supported classification and were common across individuals, group importance frequency maps indicate the number of participants for which the voxel accurately distinguished each mental state. All importance voxels were summed, irrespective of average positive or negative *z*-scored activation. Frequency maps were also computed that independently summed positive (**Fig. S2a**) and negative (**Fig. S2b**) *z*-scored activation voxels, as well as histograms of frequency counts (**Fig. S2c-e**). Note that the maximum frequency for any importance map was 10/14.

### Step 2: Decoding the focus of attention during breath meditation

Individualized brain patterns for each participant were used to decode the focus of attention during 10 minutes of breath meditation, producing a continuous estimate of internal attention states of attending to the breath, mind wandering, or self-referential processing (**Fig. 4a-d**). Classifier decisions at each time point were based on the class with the highest classifier evidence values (**Fig. S1**). From these data, “mental events” were defined whenever there were 3 or more consecutive time points that were classified as belonging to the same mental state (**Fig. 4b**). See **SI** and **Table S9** for additional analyses with alternate data reduction of classifier evidence, decisions, and mental event length.

**Figure 4.**
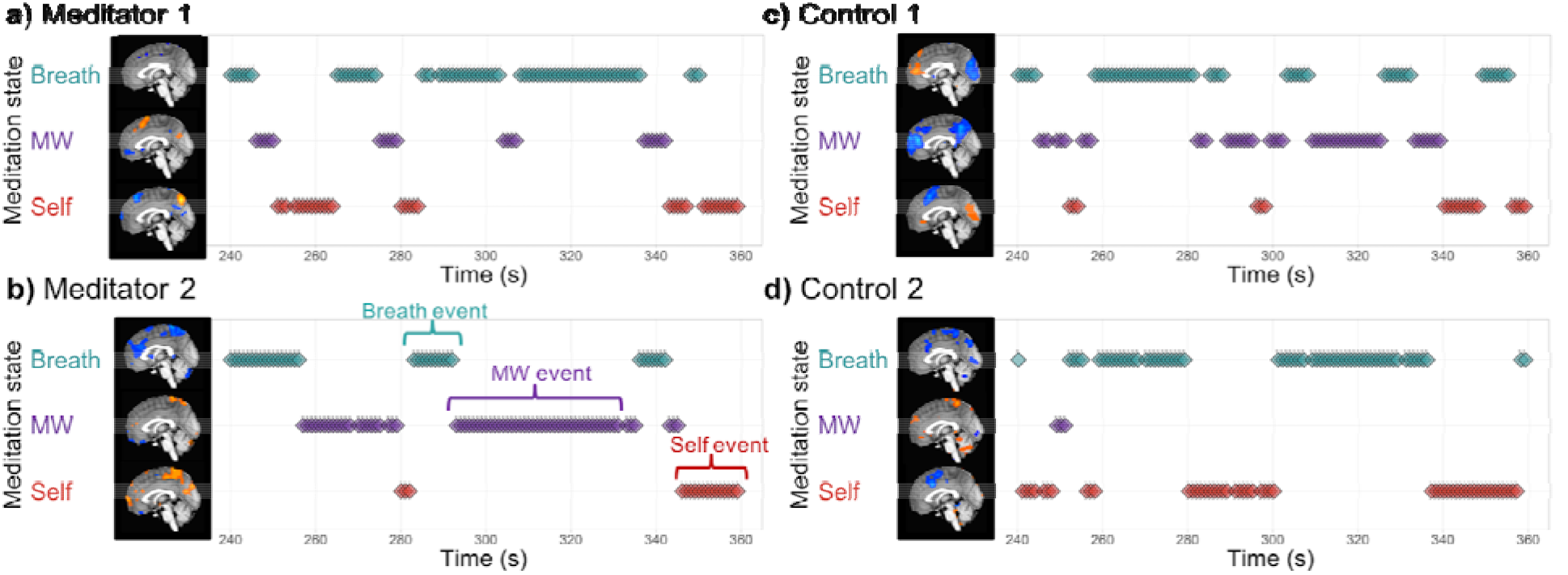
EMBODY Step 2: Decoding the internal focus of attention during breath-focused meditation using individualized brain patterns. Based on each participant’s unique brain signatures for Breath, MW, and Self, classifier decisions were made for each time point of fMRI data (TR=1s), producing a continuous estimate of attention states during breath meditation. The middle of the meditation period is displayed for two meditators **(a, b)** and their matched controls **(c, d)**. Mental events were quantified as 3 or more consecutive decisions from the same mental state **(b)**, and were used to compute metrics of attention during meditation in Step 3. See **Methods** and **SI** for details and alternate data reduction of classifier evidence and decisions.

### Step 3: Quantifying metrics of internal attention during breath meditation

Based on MVPA classification of mental states during meditation from Step 2, we computed metrics of attention during meditation for each participant, including *percentage time spent engaged* in each mental state, *number of mental events* (or discrete periods engaged in each mental state), the *duration of each mental event*, and the *variance of the durations* (SD).

#### Attention profiles during breath-focused meditation

Although the main goal of this study was to test the feasibility of using MVPA to estimate IA states during meditation at the individual level, we preliminarily characterized attention profiles at the group-averaged level (**Fig. 5**; **Table S8**). For breath-focused meditation, we hypothesized that participants would direct their attention more to the breath than engaging in mind wandering or self-referential processing. Therefore, compared to the other mental states, participants should show greater: 1) percentage time attending to the breath, 2) number of breath mental events, 3) mean duration of attention to the breath, and 4) variance in duration on the breath (greater inter-trial variability due to longer durations). Attention metrics differed in the percentage time engaged in each mental state (*F*_2,12_=8.93, *p*=0.001), the mean duration of mental events (*F*_2,12_=6.47, *p*=0.005), and the mean variance of event durations (*F*_2,12_=4.20, *p*=0.026). Consistent with our hypotheses, we found that during meditation, participants spent more time paying attention to their breath compared to mind wandering or self-referential processing (*t*_13_s>3.18, *p*s<0.007). These results remained consistent even with alternate data reduction of classifier decisions, evidence, and event length during the meditation period (see **SI** and **Table S9** for full details).

**Figure 5.**
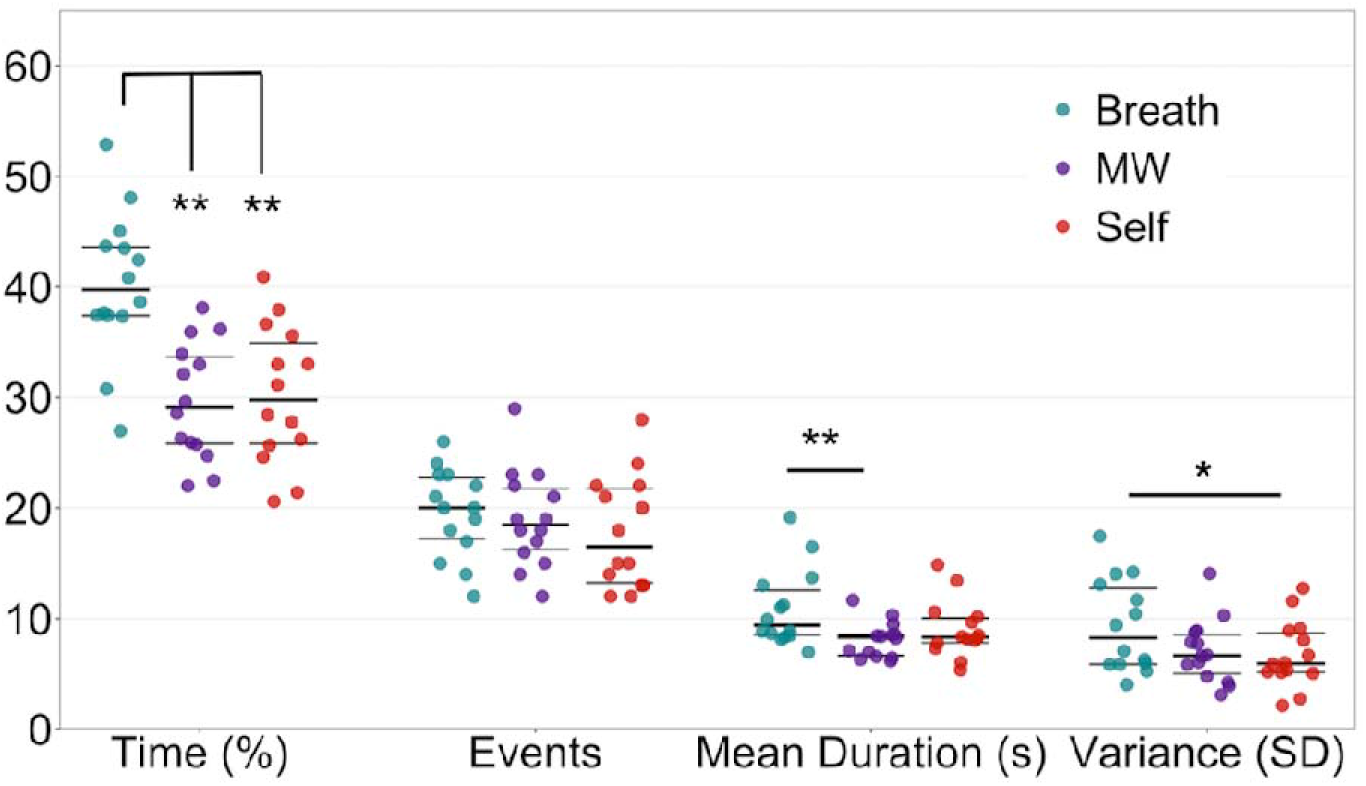
EMBODY Step 3: Quantification and mental state profiles of internal attention during meditation. Based on the estimate of mental states and event specification from Step 2, metrics of attention during breath meditation were quantified for each mental state and initially characterized at the group level: percentage time spent in each mental state (Breath, MW, or Self), the number of events, mean duration of events (s), and variability (standard deviation or SD) of duration of events. Overall, participants spent more time attending to the breath vs. mind wandering and self-referential processing. Beeswarm plots present each data point, the median (bold black line), and ±25^th^ percentile range (gray lines). See **Table S9** for full metric statistics. * paired *t*_13_=2.46, *p*=0.029, after one-way ANOVA *F*_2,12_ = 4.20, *p*=0.026 ** paired *t*s_13_≥3.18, *p*s≤0.007, after one-way ANOVA *F*s_2,12_≥6.47, *p*s≤0.005

On average, the 10-min meditation periods contained 56.4 mental events of at least 6-s each (SD=11.26). Although the mean number of events across mental states did not differ significantly (*p*=0.31), when participants attended to the breath, the mean duration of those events (10.9s [3.5]) was longer than for mind wandering events (8.1s [1.6], *t*_13_=3.28, *p*=0.006) and marginally longer than self-referential processing events (9.0s [2.6], *t*_13_=1.94, *p*=0.07). Similarly, the variability of event durations tended to be greater for attention to the breath compared to both mind wandering (*t*_13_=1.92, *p*=0.08) and self-referential processing (*t*_13_=2.46, *p*=0.029). See **Table S8** for full statistics (including distraction from breath and mental state fluctuations). See **SI** for preliminary correlations between attention metrics and between-subject variables (self-reported attention during meditation, lifetime meditation practice [**Fig. S3**] and trait interoception and mindfulness [**Table S10]**). For descriptive metrics by group, see **Fig. S4** (classifier accuracy) and **Fig. S5** (attention profiles).

## DISCUSSION

This proof-of-principle study tested the feasibility of the EMBODY framework, where MVPA pattern classifiers were applied to neural data to learn and decode internal attention states during meditation, producing novel estimates of interoceptive focus during meditation. We demonstrated that fMRI pattern classifiers could indeed distinguish between five states of internal attention using participant-specific brain patterns. This analysis was successful in all but two participants (87.5%; including all 8 experienced meditators and 6/8 novice controls), demonstrating that MVPA recognition of directed IA states has high generalizability, particularly for experienced meditators. Further, within-subject classification accuracy was positively correlated with subjective ratings of internal attention, suggesting that the EMBODY framework can reliably assess neural patterns representing internal attention. These neural patterns were then used to continuously decode the presence of breath attention, mind wandering, and self-referential processing during an independent 10-min period of breath-focused meditation. By making these invisible internal processes visible and quantifiable, we were able to compute novel profiles of attention during meditation, including the percentage of time engaged in breath attention, mind wandering, or self-referential processing (as well as number of mental events, mean duration, and variance). Preliminarily, across all participants with distinguishable brain patterns, attention profiles indicated they engaged more with the breath vs. other states (greater percentage of time attending to the breath and greater mean duration of breath events). This objective measure provides initial evidence that participants were indeed able to implement the meditative goal of sustaining attention to the breath. Together, these findings support the feasibility of employing the EMBODY framework to utilize participant-specific neural data to estimate interoceptive focus and other mental states during meditation.

To establish reliable neural patterns to decode meditation states, we first tested whether directed internal attention states could indeed be recognized by MVPA in the IA task. Interestingly, even with no changes in the external visual environment, by simply directing the internal focus of attention to five types of internal stimuli (sensations of the breath and feet, engaging in mind wandering or self-referential processing, and listening to ambient sounds), this produced reliable and distinct neural patterns for most participants. Further, initial construct validity of the IA task was supported through evidence that within-subject neural classification accuracy (which indicates reliability and distinctiveness) was positively correlated with subjective attention ratings (particularly breath trials), suggesting that more consistent IA neural patterns reflect more stable subjective IA. This approach worked in all experienced meditators and most novice controls, suggesting that the task may be applied in both cross-sectional and longitudinal study designs. However, the IA task needs further validation in larger samples, and should be improved to increase classifier performance (optimizing trial conditions and durations, testing different classification algorithms, integrating psychophysiological and behavioral data), and use real-time neurofeedback to aid interoceptive focus (Sitaram et al. 2017). These early results motivate this future work and support the feasibility of using MVPA to distinguish between different internal attention states using a participant-specific approach.

Notably, the important voxels that contributed to accurate classification for each mental state were distributed across many areas of the brain and tended to be unique for each participant, which lends support to using individualized MVPA approaches to measure IA states (see **SI** for discussion of initial group-level regions). Similar to previous research (Kerr et al. 2011), these results demonstrated that neural signals may differentiate interoception to distinct areas of the body (breath vs. feet), which could potentially track attention during body scan practices. These findings also showed that the brain patterns for mind wandering, or the “movement” from one mental state to another (Christoff et al. 2016), were distinct from self-referential processing, which demonstrated that the wandering nature of attention could be disentangled from the *contents* of what the mind wanders to.

By establishing that MVPA could recognize distinct IA states, we could then apply these classifiers to estimate the presence of IA states using neural patterns during a separate meditation period. We demonstrated the feasibility of using neural patterns to estimate mental states during meditation, producing a temporal read-out of mental states that could be used to estimate the percentage time engaged in interoceptive focus. To our knowledge, this study provided the first objective measure that enabled continuous estimation of mental states that did not impact the meditative process by requiring subjective or motor responses (Levinson *et al.* 2014). However, MVPA decisions of meditative mental states should be further validated with participant subjective reports (Hasenkamp *et al.*, 2012) and associations with other measures of interoception and mindfulness in larger samples. Although the focus of the current approach was on within-subject analyses, we preliminarily assessed differences in attention profiles across participants and found that participants spent more time attending to the breath vs. other mental states during meditation (even with alternate analyses of classifier evidence and decisions), due to longer duration on the breath when attention was directed there. This early result suggests that attention can be reliably directed towards a focused internal stimulus such as the breath; however, this should be replicated in studies with larger sample sizes, and meditative attention profiles should be compared to other states such as resting state. Additionally, given the high cost of fMRI-based measures, the framework should be extended using alternative neuroimaging methods such as electroencephalography and magnetoencephalography (Zhigalov et al. 2019).

Overall, the initial EMBODY framework shows promising ability to distinguish unique brain patterns of internal attention, which can then be used to estimate mental states during meditation. These new metrics may aid measurement of internal focus during meditation practice, which could elucidate how cultivating qualities of internal attention may transfer to cognitive and emotion regulation. Given that meditation trains multifaceted qualities of attention, the framework may be adapted to measure other aspects of attention (e.g., meta-awareness, nonjudgment) and meditation practices (e.g., open monitoring, compassion). By developing measures to precisely assess the attentional qualities cultivated by meditation, we will gain the measurement power needed to rigorously test the attentional and emotional mechanisms through which meditation may improve health and well-being. Finally, the EMBODY framework highlights that each individual’s brain signatures and meditation practice are unique, which we hope will aid researchers and clinicians in designing interventions that will maximally benefit individuals in targeted and specific ways.

## ACKNOWLEDGEMENTS

This work was supported by the US National Institutes of Health [grant numbers K08 AT009385 [H.Y.W.], T32 AT006956 [F.M.H. and S. Adler], K24 AT007827 [F.M.H.], R01 EY028746 [J.A.L.P], R01 AG049424 and R21 AG041071 [D.A.Z. and A.G.], the UCSF Mt. Zion Health Fund Pilot in Integrative Medicine Research [H.Y.W.], and Osher Center for Integrative Medicine (OCIM) Jaswa Fund for Meditation Research and Bowes Foundation Research Fund [H.Y.W.]. This manuscript has been released as a pre-print at bioRxiv (Weng et al. 2018). The authors thank Lara Stables, Chad Smiddy, Elizabeth Pierce, Peter Wais, Ryan Lopilato, and Sierra Niblett (UCSF Neuroscape) for MRI consultation and sequences, study management, and data collection; Patricia Moran and Stephanie Lee (UCSF OCIM) for study management, screening, and data collection; Mark Estefanos for aid in software development; and Tiffany Ho and Regina Lapate for consultation regarding study design, data analysis and interpretation.

## AUTHOR CONTRIBUTIONS

H.Y.W., J.L.P, F.M.H., and N.A.S.F. conceptualized the study. All authors contributed to the data analytic strategy and interpretation. H.Y.W., S.S., and V.G. contributed to data collection, processing, and analysis. H.Y.W., J.L.P., S.S., and V.G. developed unpublished data analysis tools. H.Y.W. wrote the manuscript, with contributions from J.L.P. and N.A.S.F. and comments from all other authors.

## Data availability and computer code

MRI data will be available for participants who consented to share raw brain data at neurovault.org (accession codes will be available before publication). Code for the EMBODY Task, MVPA analysis, and post-processing are available upon request.

