## Supplemental Information for "Focus on the breath: Brain decoding reveals internal states of attention during meditation"

**Table of Contents**

| **Supplemental Methods and Results** | Page |
| --- | --- |
| **EMBODY Step 1 – Internal Attention (IA) Task** |  |
| Classifier confusion matrix | 2 |
| Classifier accuracy by group | 2 |
| Influence of head motion | 2 |
| Influence of respiration | 3 |
| Common brain regions contributing to accurate IA classification | 4 |
| **EMBODY Step 2 – Decoding mental states and data reduction** |  |
| Alternate analyses of classifier decisions and evidence | 5 |
| **EMBODY Step 3 – Meditation Period Metrics** |  |
| Additional metrics: distraction from breath and mental state fluctuations | 7 |
| Exploratory group differences in meditation metrics | 7 |
| **Exploratory construct validity with meditation metrics** |  |
| Meditation period ratings | 8 |
| Lifetime meditation practice | 8 |
| Trait interoception and mindfulness | 9 |
| **Supplemental Figures** |  |
| **Figure S1.** EMBODY meditation decoding output: classifier decisions and evidence | 11 |
| **Figure S2.** Frequency count of positive and negative importance voxels | 12 |
| **Figure S3.** Exploratory associations of mental states during meditation and lifetime  meditation practice | 13 |
| **Figure S4.** Classifier accuracy by group | 14 |
| **Figure S5.** Exploratory group differences in EMBODY meditation period metrics | 15 |
| **Supplemental Tables** |  |
| **Table S1.** Demographic information | 16 |
| **Table S2.** Meditation practice traditions and statistics | 17 |
| **Table S3.** Head motion by Internal Attention task condition | 18 |
| **Table S4.** Respiration rate in the IA task | 18 |
| **Table S5.** Internal Attention (IA) task classifier confusion matrix | 19 |
| **Table S6.** Individual-level classification accuracies from the IA task | 19 |
| **Table S7.** Brain regions from group frequency importance maps, see TableS7.xlsx | 20 |
| **Table S8.** Preliminary group-level meditation period metrics | 20 |
| **Table S9.** Meditation metrics from alternate analyses of meditation period | 21 |
| **Table S10.** Exploratory associations between meditation metrics and trait  interoception and mindfulness | 22 |
| **Supplemental References** | 23 |

**Supplemental Methods and Results**

**EMBODY Step 1 – Internal Attention (IA) Task**

**Classifier confusion matrix.** The classifier confusion matrix consists of the number of classifier decisions for each category given the instructed condition in the IA task (**Table S5**), averaged across all 16 participants. 432 total decisions were made for each second of data in each category in the IA task (see Online Methods for full classification details using MVPA(Norman *et al.*, 2006)). In each condition, the greatest number of classifier decisions were made for the correct congruent category (**Table S5**), indicating that the classifier could accurately recognize the brain pattern for each condition. Breath brain patterns were most likely to be confused with Feet brain patterns, indicating confusion with attention to another area of the body. Mind Wandering (MW) brain patterns were most likely to be confused with self-referential processing (Self) brain patterns, whereas Self brain patterns were most likely to be confused with Feet brain patterns. Finally, the control conditions of Feet and Sounds were most likely to be confused with each other. Mean classifier accuracies and SD of each condition are also listed in **Table S5** to aid future research (displayed in **Fig. 3** in the main manuscript).

**Classifier accuracy by group.** Mean classifier accuracies from the IA task were computed for each group (8 Meditators and 8 Controls) for exploratory purposes and to aid future research. Within each group, accuracy for each condition was tested with a one-sample *t*-test vs. 20% (theoretical chance level for 5 conditions; **Fig. S4**). Differences between groups were tested with a Mann-Whitney test in each condition. Each condition was recognized significantly above chance in each group, including the three conditions of most interest for breath-focused meditation (Breath, MW, Self) and the two control conditions (Feet, Sounds). The groups did not differ in classifier accuracy in any condition (*p*s > 0.19).

**Influence of head motion.** To investigate the potential influence of head motion(Churchill *et al.*, 2012), we compared motion in each of 5 conditions in the IA task (Breath, Feet, MW, Self, and Sounds). Using the 1D files output by AFNI’s 3dvolreg,we computed mean motion in each of 6 directions (roll, pitch, yaw, superior-inferior [dS], left-right [dL], posterior-anterior [dP]) as well as total mean motion across all 6 directions (7 metrics total). Difference in motion between all 5 conditions were tested using a one-way ANOVA for each direction and total motion. ANOVAs were also computed for the 3 main conditions of interest only (Breath, MW, Self). Head motion did not vary between the 5 conditions in any of 7 motion metrics (all one-way ANOVA *F*s_4,75_<1.35, *p*s>0.25; **Table S3**), nor between the 3 conditions relevant for meditation (Breath, MW, Self; all *F*s_2,45_<0.73, *p*s≥0.48).

Although head motion did not vary by condition, to ensure classifier accuracy was not substantially driven by motion, we re-calculated IA task classification accuracy with fMRI data where motion was covaried out of the BOLD signal using 3dDetrend. Classifier accuracy remained robust for each condition (all mean accuracies>39% vs. 20% chance, all *t*_15_s>6.03, all *p*s<0.0001). We therefore did not correct for head motion in the main analysis. However, to mitigate the impact of individual-level head motion on identifying voxels in the importance map analyses, we conducted the importance map analysis on fMRI data with head motion covaried out (**Figs. 3** and **4** in main manuscript**, Fig. S2**).

**Influence of respiration.** Previous research demonstrates respiration rate may change depending on meditative state and practice experience(Farb *et al.*, 2013; Wielgosz *et al.*, 2016), which may subsequently influence blood-oxygen-level dependent (BOLD) fMRI activity. Before fully discussing the influence of respiration in this experiment, it is important first to clarify the conceptual and methodological differences between standard univariate approaches and our novel multivariate approach (MVPA) in analyzing fMRI data to study meditation. In the standard univariate approach, brain data from different cognitive states (e.g., meditation vs. rest) are compared and averaged for the purpose of *brain mapping*, or identifying brain regions associated with mental states at the group level. In this case, the location in the brain is the main outcome, and therefore it would be important to regress out activation due to non-neural sources such as respiration. However, in our multivariate approach with the EMBODY framework, we are instead using the brain data at the individual level to *distinguish and decode cognitive states* during meditation(Norman *et al.*, 2006). The resulting *cognitive states are the main outcome of interest* rather than brain activity *per se*. Therefore, any source of information that contributes to differential brain patterns in the BOLD signal are useful (such as physiology), and should *not* be regressed out as they would remove important diagnostic signals.

We therefore frame our investigation of the influence of respiration as secondary analyses to understand whether 1) respiration rate changes due to IA task instruction (particularly for Breath, MW, and Self), and 2) accurate classification accuracy in the IA task is primarily driven by changes in respiration signal.

**Methods.** Respiration data were collected using a Siemens respiration belt at a sampling rate of 51 Hz. Full respiration datasets were available for only 3/16 participants due to technical errors (such as faulty respiration belt signal). One participant was an experienced meditators, and two were control participants (one of which did not show accurate classification accuracy in the main analysis pipeline). With this subsample, we examined the influence of internal attention on respiration rate (RR) at the trial level within each subject using in-house Python software. The raw respiration data were linearly detrended and processed with a low-pass filter (5^th^-order Butterworth filter) with a cut-off frequency of 0.5 Hz. Peaks and troughs were identified as local maxima and minima within the respiratory signal. Respiratory phase was modeled by coding peaks (1), troughs (-1), and neither peak nor trough (0). The number of breaths were counted per trial as trough-to-trough intervals, and converted to respiration rate (RR) in breaths per minute for the main conditions of interest: Breath, MW, and Self.

**Results.** Mean RR between each pair of conditions was tested with paired *t*-tests using the first 12 trials in each condition (more breath trials were administered at shorter durations). Because this analysis is exploratory to examine whether internal attention influences RR, we did not correct for multiple comparisons and report all tests for completeness and to encourage further research. Each participant showed slower RR for Breath vs. MW (all *t*s<-3.31, all *p*s<0.01), and 2/3 participants showed slower RR for Breath vs. Self (*t*s<-3.40, *p*s<0.01). No participants showed significant differences in RR for MW vs. Self. Subject-level data are reported in **Table S4.**

Because differences in RR between conditions were found, we regressed the processed respiration waveform from the fMRI BOLD data using AFNI’s 3dRetroIcor (before slice-time correction, as recommended by AFNI), and found that Step 1 Internal Attention task classifier accuracy remained similar to the primary analysis without respiration covaried out (mean difference = 0.60%, range of -4.17% to 3.94%). Individual-level classification accuracy was tested using Chi-Square tests, and yielded the same significance result in each condition in each subject as the primary analysis (2 subjects showed significant accuracy in all 3 conditions, 1 subject did not have accurate classification in any condition). For this small sample, the results suggest that classifier accuracy is *not* primarily driven by variations in respiratory signals. However, future studies should investigate these questions more rigorously, and investigate how decoded mental state transitions during meditation relate to temporal parameters of physiology.

**Common brain regions contributing to accurate IA classification.** Voxels that were important for breath-focused attention in a higher frequency of participants (N≥5) were located in the left lateral temporal cortex (spanning the superior temporal gyrus, middle temporal gyrus, and middle frontal gyrus), left inferior temporal gyrus, bilateral precuneus, and right frontal pole. Higher-frequency importance voxels for mind wandering (N≥5) were located in bilateral precuneus, bilateral anterior medial prefrontal cortex (PFC), left precentral and postcentral gyrus, and other areas of the PFC (bilateral frontal pole, right middle frontal gyrus). Higher-frequency importance voxels for self-referential processing (N≥6) were located in bilateral dorsomedial PFC, bilateral precentral gyrus, left superior parietal lobule, right anterior insula/frontal operculum cortex, and areas of the PFC (right frontal pole, dorsolateral, middle frontal gyrus) (**Fig. 3b**, **Fig. S2**, **Table S7**).

Although this framework emphasizes the individual differences in brain patterns for person-specific decoding of meditation, we initially characterized brain regions that may contribute to accurate classification in a higher proportion of participants. Note that regions identified by these group-level importance maps highlight voxels that contribute to accurate classification between conditions, and do not depict the same neural metrics as standard univariate analyses (e.g., comparing mental states to find differences in average activation). For breath-focused attention, clusters spanned the lateral temporal cortex (LTC, including areas in the superior and middle temporal gyrus, and middle frontal gyrus) and inferior temporal gyrus, and consisted mostly of negative importance voxels, indicating that the average *z*-scored activation was lower than mind wandering and self-referential processing. The neural signature representing breath attention may involve the relative de-activation of temporal regions due to their association with the dorsal medial prefrontal cortex (mPFC) subsystem of the Default Mode Network (DMN), which includes the lateral temporal cortex and temporal pole, and is thought to be involved in making judgments about present mental states (Andrews-Hanna *et al.*, 2010). Surprisingly, we did not find a consistent cluster in the insula which is important for interoception (Craig, 2009), and this null result may stem from the lack of an external stimulus (often used as a comparison condition; Farb *et al.*, 2013)). However, self-referential processing was characterized by a negative importance cluster in the anterior insula/frontal operculum cortex, which suggests it may be identified in part by less activation in regions typically associated with interoception.

Self-referential processing also involved a large negative importance cluster in the dorsomedial PFC, which is typically active in the task-positive Executive Function Network (EFN; Fox *et al.*, 2016). Finally, prevalent regions for mind wandering included the anterior mPFC and bilateral precuneus (adjacent to the posterior cingulate cortex) and consisted of negative importance voxels. The anterior mPFC and posterior cingulate cortex are key hubs of the DMN midline core that reflect self-relevant affective decisions (Andrews-Hanna *et al.*, 2010), and thus less DMN activation may represent an important signal in distinguishing mind wandering vs. self-referential processing. Surprisingly, few regions with positive importance voxels emerged in the analyses, which suggests that predictive voxels indicating positive average activation within a given condition may be more distributed throughout the brain with less overlap between individuals. Research with larger samples may further characterize group-level neural patterns that can be used to decode meditation, and different analyses of the fMRI signal may indicate which predictive voxels are associated with physiological variables such as respiration and heart rate.

**EMBODY Step 2 – Decoding mental states and data reduction**

**Alternate analyses of classifier decisions and evidence.** The default setting in MVPA Toolbox is to base categorical classifier decisions off of the condition assigned the highest continuous classifier evidence value (a “winner-take-all” approach; **Fig. S1**). This may obscure data points that represent transitions between mental states, or be incongruent with current theories which allow for mind wandering to occur simultaneously with on-task performance(Thomson *et al.*, 2015). Therefore, the classifier evidence and subsequent decisions can be parsed in many additional ways that represent more nuanced conceptualizations of fluctuating mental states. Classifier evidence can be **a)** analyzed as *overall probabilistic values* across the entire meditation period, or **b)** contribute to *more stringent decisions which require a minimum distinction* between the two highest evidence values (e.g., requiring a 0.2 or 20% difference in evidence values between the top two categories, say Breath and MW). This method could discard volumes that represent a mixed transitional state between mental states, rather than assigning them a category. Further, once decisions are made to identify mental states during meditation, data can be cleaned by **c)** requiring a *minimum “mental event” length* (main analysis: 3s) to identify more stable mental states, and **d)** implementing a *smoothing algorithm* which re-labels a single incongruous decision between two events of the same type (e.g., MW event – Self decision – MW event) according to the category of the surrounding events (e.g., Self => MW; main analysis smoothed an average of 1.3% data points, SD = 0.41).

In the main analysis reported in the manuscript, we used the default MVPA decision setting (highest evidence value), applied a smoothing algorithm, and required a mental event to be ≥3s long. To examine the consistency of our main findings where percentage time engaging in the breath was greater than engaging in mind wandering or self-referential processing, we also re-analyzed the meditation period evidence data as overall probabilistic values across all data points, as well as based on a range of minimum evidence distinction requirements (0.05-0.30 or 5-30% distinction). Classifier decision data were also re-analyzed based on minimum evidence distinctions, with smoothed or unsmoothed data, and varying mental event lengths (2, 3, or 4s). Paired *t*-tests of percentage time engaging in Breath vs. MW and Breath vs. Self were then re-computed (**Table S9**).

a. **Overall probabilistic values during the meditation period.** This analysis computed the mean evidence values for Breath, MW, and Self for each participant across the entire meditation period. When this was done for all data (with no exclusion parameters), mean evidence for Breath was greater than MW and Self (Breath=0.51, MW=0.49, Self = 0.49; *p*s<0.05). We also computed this metric for each parameter of excluding ambiguous frames from the previous analysis (evidence difference of 0.05 – 0.3 (or 5-30% difference) in 0.05 increments. This analysis required that for each TR, the condition with the highest evidence value had to be at least 0.05 – 0.3 units greater than the second-highest condition’s evidence value. Each analysis produced a new read-out of classifier data points, and ranged in the amount of data points excluded (17.5%-48.6%). Evidence probability of engaging in each mental state across the entire meditation period was re-computed based on the included data points. For each of these 6 analyses, we still found that the evidence probability engaged in Breath during meditation was greater than MW or Self (all *p*s<0.05), and there were no differences between MW and Self. Generally, this finding became stronger as the evidence distinction level became higher.
b. **Classifying mental states by excluding ambiguous frames.** We excluded ambiguous data points based on minimum distinction criteria between evidence values (range 0-1). For each volume of data during the meditation period, we required a minimum distinction (or difference) between the two conditions with the highest evidence values, ranging from 0.05–0.3 (or 5-30% evidence difference) in 0.05 increments, resulting in 6 total distinction levels. Each analysis produced a new read-out of classifier decisions, and excluded frames that did not meet the criterion. The percentage of data excluded ranged from 17.6% (0.05 distinction) to 48.6% (0.3 distinction). The main finding remained significant for all 6 analyses, where the percentage time engaged in breath attention was greater than MW or Self (all *p*s<0.01).
c. **Variable “mental event” length and d. implementing a smoothing algorithm.** To identify more stable mental events, in the main analysis, we required mental events to be at least 3s long and implemented a smoothing algorithm which re-labels a single incongruous decision between two events of the same. To investigate the consistency of the main group-level finding that participants spent more time attending to the breath vs. engaging in mind wandering or self-referential processing, we re-computed these tests for all alternate analyses of the meditation period (data that were raw, unsmoothed or smoothed, event length ≥2, 3, or 4s). Using paired *t*-tests, we found that the main findings were stable for all 7 analyses (Breath > MW, all *t*s_1,13_>3.55, all *p*s<0.005; Breath > Self, all *t*s_1,13_>3.03, all *p*s<0.01; **Table S9**).

Across all participants and analyses, percentage values did not greatly differ from the main analysis (smoothed, event length ≥3s) for Breath (mean difference=-0.28%, SD=1.63), MW (mean difference=0.02%, SD=0.53), or Self mean difference=-0.15%, SD=0.54). This suggests that slight variations in the way the data were parsed did not greatly change the overall percentage time metrics. However, other metrics such as mean duration were sensitive to these slight changes (i.e., increasing minimum event length from 2 to 4s would increase the mean duration of events), so we present these findings in **Table S9** to inform analysis of future datasets. Although mean duration lengths changed depending on analysis, the main finding that the mean duration of Breath was greater than MW remained significant for each analysis (all *t*s_1,13_>2.92, all *p*s<0.05), and the mean duration of Breath was greater than Self remained significant for 4/6 analyses (*t*s_1,13_>2.22, all *p*s<0.05).

Analyses greatly ranged in the amount of data excluded, from 7.57% (smoothed, event length≥2) to 23.58% (unsmoothed, event length≥4). Because these main percentage time results were stable across all analyses, we chose to present our main analysis in the paper (smoothed data, event length≥3s) which has a moderate amount of data excluded (15.73% data), and reported the other analyses (**Table S9).** Based on these analyses, we are more confident in the robustness of the main finding that participants are indeed able to pay attention to their breath longer than engaging in mind wandering or self-referential processing during breath-focused meditation.

**EMBODY Step 3 – Meditation Period**

**Additional metrics: Distraction from breath and mental state fluctuations.** From the classifier decisions output from Step 2, the total number of mental events and mean duration of distraction from breath were also calculated. Fluctuations between mental states were quantified by counting the number of transitions from each mental event (Breath, MW, Self) to the next mental event (Breath, MW, Self). For each mental state type, we tested the difference in the mean counts of subsequent transitions to the other mental state types (e.g., after Breath events, the difference in mean count between transitions to MW and Self), using pair-wise *t*-tests. When participants became distracted from their breath, the distraction period lasted for an average of 20.4s (SD=4.77), and was marginally more likely to be attributed to entering a state of mind wandering (mean count=8.8 [2.46]) than self-referential processing (mean count=7.0 [2.57]; *t*_1,13_=1.90, *p*=0.08). When engaged in mind wandering, participants were equally likely to transition to mental states of breath (mean count=8.6 [2.77]), or self-referential processing (mean count=7.9 [3.85], *p*=0.68), and when engaged in self-referential processing, they were equally likely to transition to breath (mean count=7.5 [2.85], or mind wandering (mean count=7.4 [2.90], *p*=0.92).

**Exploratory group differences in meditation metrics.** Although the sample size is not large enough to adequately explore differences between meditators and controls, we performed exploratory group tests in meditation metrics from the EMBODY Task. We hope this will help generate hypotheses for future research. Results should be interpreted with caution due to small sample size, and future research with larger sample sizes should extend these findings. With a larger sample size, we hypothesize meditators would focus on their breath longer, and spend less time engaged in mind wandering and self-referential processing.

We first determined whether each metric varied by group and condition (Group * Condition repeated measures ANOVA, performed on ranked values to account for small sample sizes). We interrogated any significant or trend-level finding for exploratory purposes only and to inform future research. We found a significant Group * Condition interaction for number of mental events (*F*_2,12_ = 4.12, *p* < 0.05), a trend-level interaction for percentage time engaged in mental events (*F_2_*_,12_ = 2.83, *p* = 0.08), and no interaction for mean duration or variance of events (*p*s > 0.36) (**Fig. S5**). Exploratory post-hoc testing for the number of events suggested that near trend-level, meditators show more instances of engaging in self-referential processing during meditation than novice controls (*t*_2,12_ = 1.74, *p* = 0.11; no difference in Breath or MW, *p*s > 0.38). Although the Group * Condition interaction for percentage time was trend-level, we performed exploratory post-hoc testing to aid future research questions. This suggested that meditators spent more percentage time engaged in self-referential processing during meditation than controls (*t*_2,12_ = 2.66, *p* < 0.05; no difference in Breath or MW, *p*s > 0.21). No group differences were found in average time distracted from breath (*p* = 0.72). All results are preliminary and should be interpreted with caution. Future research should include larger sample sizes to make inferences about the differences between meditators and controls.

**Exploratory construct validity with meditation metrics**

To conduct exploratory examinations of construct validity, EMBODY Task metrics were correlated with subjective measures of meditation period ratings, lifetime meditation hours, and trait questionnaires of interoception and mindfulness. Given the small sample size to conduct between-subjects analyses (N=14 or 8), these results are reported as descriptive statistics for exploratory purposes and to inform future research.

**Meditation period ratings**. After the meditation period, participants rated the percentage time they paid attention to their breath and to their thoughts, and these ratings were not correlated with percentage time metrics (breath *rho*_13_=-0.05; self *rho*_13_=-0.09).

**Lifetime meditation practice.** During the phone interview, amount and type of meditation practice was assessed. Participants reported total years of consistent meditation practice (defined as $\geq$90 minutes a week). If there was a break in practice, each period was assessed independently. For each time period of consistent practice, both total weekly group practice and personal practice were reported. Participants reported total years of practice and average minutes per week of each type of practice. Participants reported what percentage of their weekly practice involved attention to breath sensations, body sensations (not primarily focused on the breath), and other practices not primarily focused on the breath or body (such as lovingkindness and mantra practice). Amount and type of monastic practice was also assessed. Participants reported months of monastic practice, formal meditation practice per day, and percentage time attending to breath, body, and other practices. Finally, participants reported history of retreat practice in 5-year blocks. For each block of time, they reported total days of practice, average formal meditation hours per day, and percentage of practice focused on breath, body, and other practice. Experience with other mind-body practices such as yoga and Tai Chi was also assessed. Based on these reports, total lifetime hours of practice were computed for all meditation practices, and meditation focused primarily on the breath, the body, and other practices. For exploratory purposes, we correlated lifetime meditation hours with EMBODY meditation metrics using non-parametric Spearman’s *rho* for descriptive purposes only. Future research should test these associations with larger sample sizes.

Within meditators, total reported lifetime meditation hours were positively associated with percentage time spent attending to the breath (*rho*_7_=0.71; **Fig. S3a**) and negatively associated with percentage time engaged in self-referential processing (*rho*_7_=-0.71; **Fig. S3b**) during the meditation period. Lifetime practice and percentage time engaged in mind wandering were not highly associated (*rho*_7_=-0.17, *p*=0.69). Furthermore, suggesting specificity of the task metrics, the amount of lifetime hours meditating *specifically* on breath sensations was positively associated with percentage time attending to the breath during the meditation session (*rho*_7_=0.74; **Fig. S3c**) and negatively associated with self-referential processing (*rho*_7_=-0.91; **Fig. S3d**). Lifetime hours meditating on other body sensations or other meditation practices were not strongly associated with any mental state during meditation (**Fig. S3e-f**).

**Trait interoception and mindfulness – *a priori* hypotheses and exploratory correlations.** To test construct validity of EMBODY attention metrics during the meditation period, we correlated them with trait-level questionnaires of interoception and mindfulness. We should note that in general, our confidence that trait-level subjective measures would correlate with EMBODY metrics is relatively low. These scales assess participants’ tendencies to sustain attention in general, and were not designed to specifically measure attention to the body during meditation practice. In addition, they depend on accurate introspection, which may be difficult given that people tend to mind wander around 30-50% of the time (Smallwood and Schooler, 2015), and may vary in meta-awareness of mind wandering or estimate the amount of time focused attention or mind wandering occurs.

Trait interoception and mindfulness were measured with the Multidimensional Assessment of Interoceptive Awareness (MAIA; Mehling *et al.*, 2012) and Five Facet Mindfulness Questionnaire (FFMQ; Baer *et al.,* 2006), respectively. Our *a priori* hypotheses were strongest for subscales that assessed sustained attention, particularly to the body. These included the **MAIA Noticing** and **Attention Regulation** subscales (ranges from 0 [never] to 5 [always])**,** where the Noticing scale assessed the awareness of uncomfortable, comfortable, or neutral body sensations (example item: “I notice changes in my breathing, such as whether it slows down or speeds up”), and the Attention Regulation scale assessed the ability to sustain and control attention to body sensations (example item: “I can pay attention to my breath without being distracted by things happening around me”). The **FFMQ** subscales included **Observing** and **Acting with Awareness** (ranges from 1 [never or rarely true] to 5 [very often or always true]), where Observing assessed noticing and attending to sensations, perceptions, thoughts, and feelings (example item: “I pay attention to sensations, such as the wind in my hair or sun on my face”), and Acting with Awareness assessed awareness of acting on habitual “automatic pilot”, and the ability to concentrate and not become distracted (example item: “When I do things, my mind wanders off and I’m easily distracted [reverse-scored]”). Note that the original FFMQ was mistakenly administered with only 14/39 total items and was not used for analyses. Full FFMQ data were collected for 13/16 participants after recognizing the error, ranging from 2-18 months (mean = 12) after the brain scan. 11 participants had valid EMBODY metrics and full FFMQ scores that could be used for correlation tests, and these data should be interpreted with caution.

Scale scores were correlated with EMBODY metrics, particularly percentage time attending to the breath, for which we had the strongest hypotheses. We used Spearman’s *rho* to account for the low sample size (N=14), and report statistics for descriptive purposes only to inform future research. Out of the four *a priori* subscales, all showed a negative association with percentage time attending to the breath, with the correlation being stronger for Attention Regulation (*rho*_13_ = -0.58; see **Table S10** for correlations with all subscales and mental states). For non-breath mental states, the FFMQ Observing subscale showed a moderate positive association with percentage time mind wandering, and the MAIA Attention Regulation subscale showed a moderate positive association with percentage time engaging in self-referential processing. These findings were not in the direction expected, and may indicate inaccurate subjective reporting, or a mismatched construct of what the trait questionnaire and meditation period metrics are measuring.

All other subscales of the MAIA and FFMQ were correlated with EMBODY metrics for exploratory purposes only (see **Table S10**). Generally, these subscales measure aspects of mindfulness and interoception that are higher-level processes that build upon more basic skills like sustaining attention to the body (e.g., self-regulation), or assess nonjudgment which the current EMBODY task did not measure. The other scales of the MAIA include: **Not Distracting** which assesses the tendency not to use distraction to cope with discomfort (“I distract myself from sensations of discomfort”), **Not Worrying** which assesses the tendency not to experience emotional distress with physical discomfort (“I can notice an unpleasant body sensation without worrying about it”), **Emotional Awareness** which assesses awareness of the connection between body sensations and emotional states (“I notice that my breathing becomes free and easy when I feel comfortable”), **Self-Regulation** which assesses the ability to regulate distress by attention to body sensations (“When I bring awareness to my body I feel a sense of calm”), **Body Listening** which assesses the tendency to actively listen to the body for insight (“I listen for information from my body about my emotional state”), and **Trusting** which assesses the experience of one’s body as safe and trustworthy (“I trust my body sensations”). The other subscales of the FFMQ include: **Describe** which assesses describing and labeling with words (“I’m good at finding words to describe my feelings”), **Nonjudge** which assesses nonjudging of experience (“I make judgments about whether my thoughts are good or bad”), and N**onreact** which assseses nonreactivity to inner experience (“When I have distressing thoughts or images, I just notice them and let them go”).

All exploratory subscales showed a negative association with percentage time attending to breath, with MAIA Not Distracting showing a stronger association (*rho*_14_ = -0.63). Several subscales also showed a positive association with percentage time engaging in self-referential processing during breath-focused meditation, including MAIA Not Distracting (*rho*_13_ = 0.73) and Body Listening (*rho*_13_ = 0.54). Future research should investigate these associations with a larger sample.

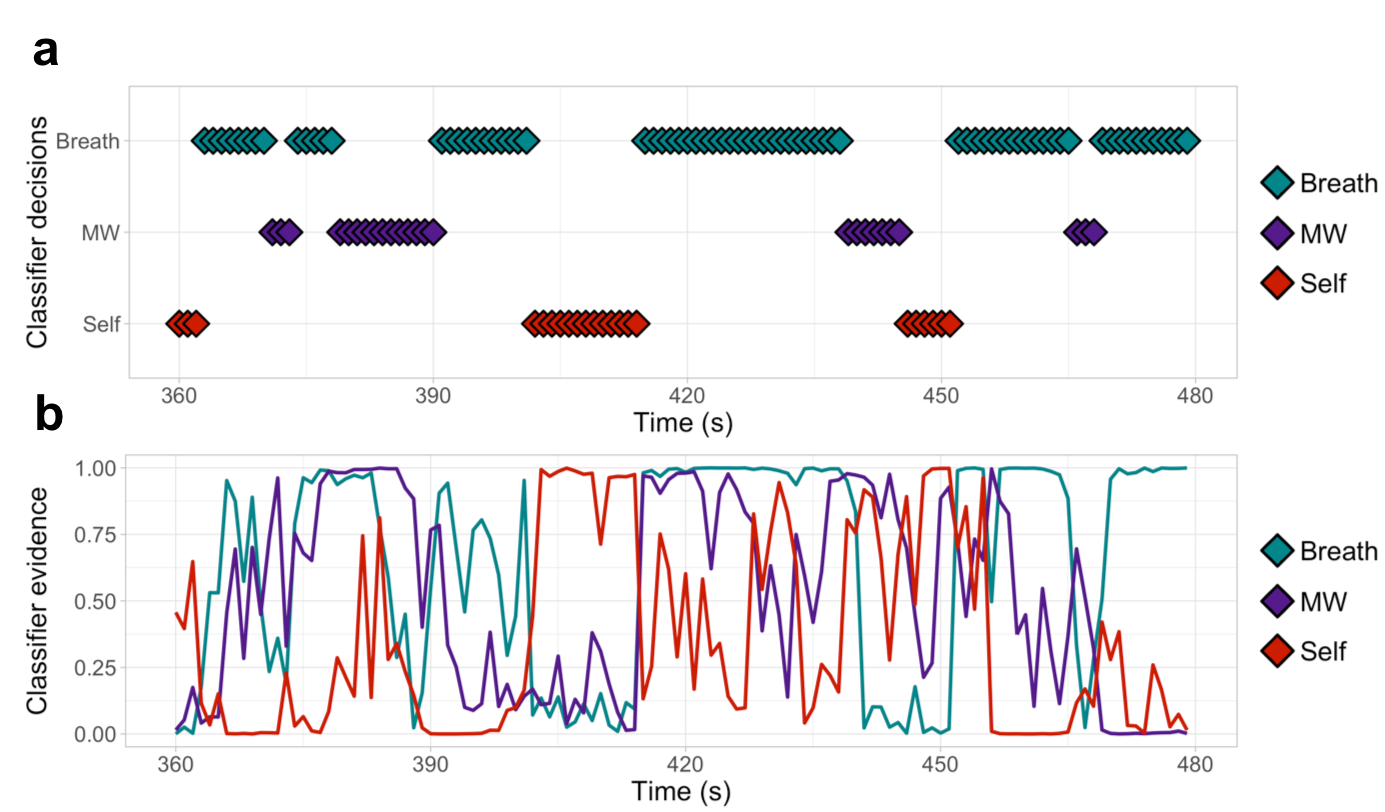

**Figure S1.** EMBODY Step 3 meditation decoding output: classifier decisions and evidence. Based on unique brain patterns learned from the Internal Attention task (Step 1), internal mental states during breath meditation are decoded by MVPA(Norman *et al.*, 2006) (TR=1s ). Outputs include **a)** categorical classifier decisions (Breath, MW, or Self) and **b)** continuous classifier evidence values (0-1) for each mental state. Classifier evidence indicates how closely each brain pattern during meditation matches the trained brain patterns (from Step 1) for each mental state. For each brain pattern, the classifier decision is based on the mental state that holds the highest evidence value. Note that classifier evidence can represent the waxing and waning of multiple mental states through time. See **SI-Methods** and **Table S7** for alternate data reduction of classifier evidence and decisions.

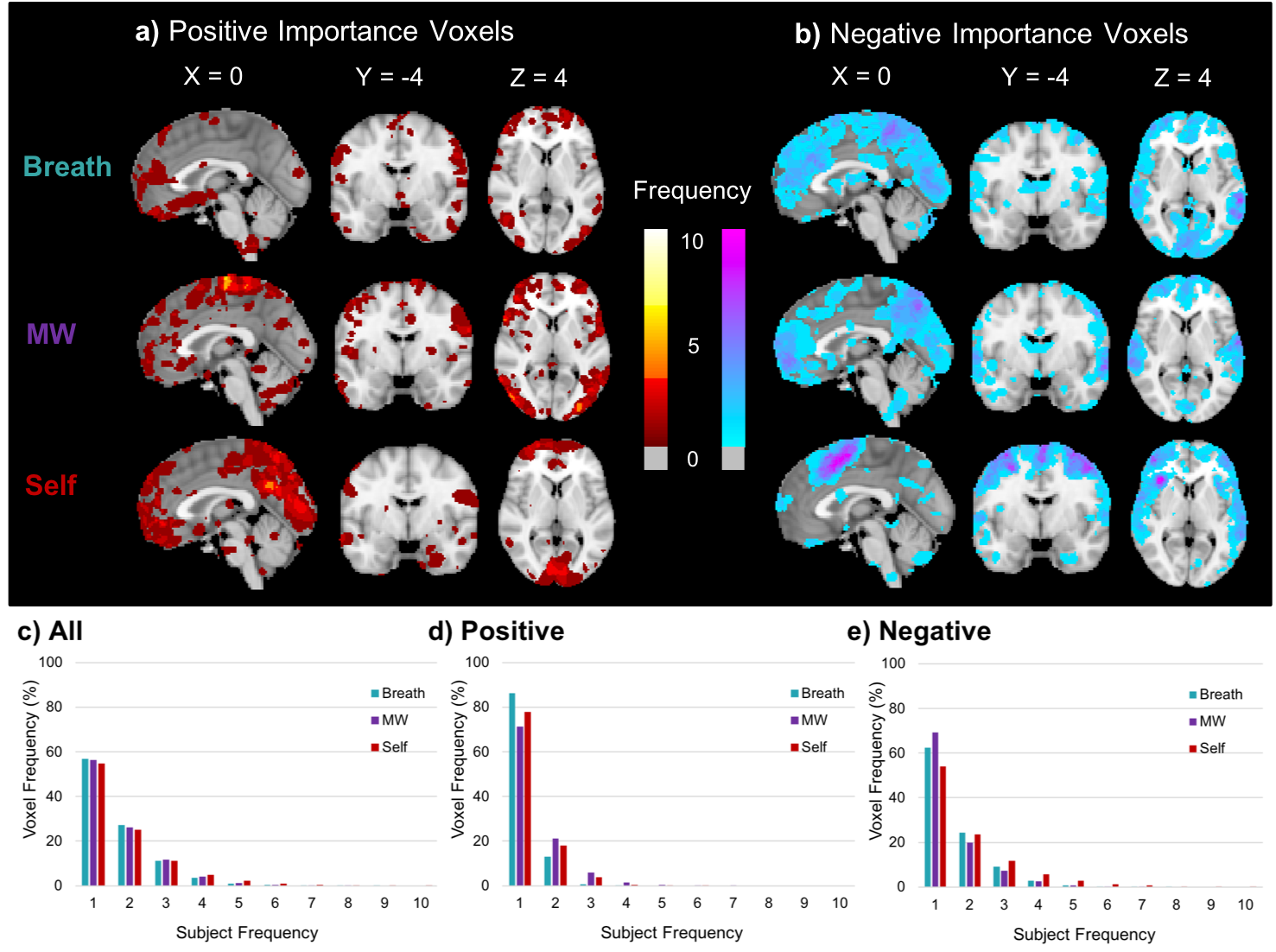

**Figure S2.** Frequency count of positive and negative importance voxels, identified using fMRI data with the influence of head motion removed. **Fig. 3b** from the main manuscript displays the frequency map of all importance voxels across 14 participants, regardless of the direction of average *z*-scored activation of the voxels. This figure displays the frequency maps specific for **a)** “positive importance” voxels, which have a positive weight and positive *z*-scored average activation value (indicating that it was more active on average, displayed in warm colors), and **b)** “negative importance” voxels, which had a negative weight and a negative *z*-score average activation value (indicating that it was less active on average, displayed in cool colors)(McDuff *et al.*, 2009). The frequency indicates the number of subjects for which that voxel was important in distinguishing the mental state (Positive maximum for Breath = 4/14, MW = 8/14, Self = 10/14; Negative maximum for Breath = 8/14, MW = 7/14, Self = 10/14). Histograms are displayed for the percentage of voxels at each subject frequency (1-10) for **c)** all, **d)** positive, and **e)** negative importance voxels. Regions with higher frequencies are reported in **Table S5**.

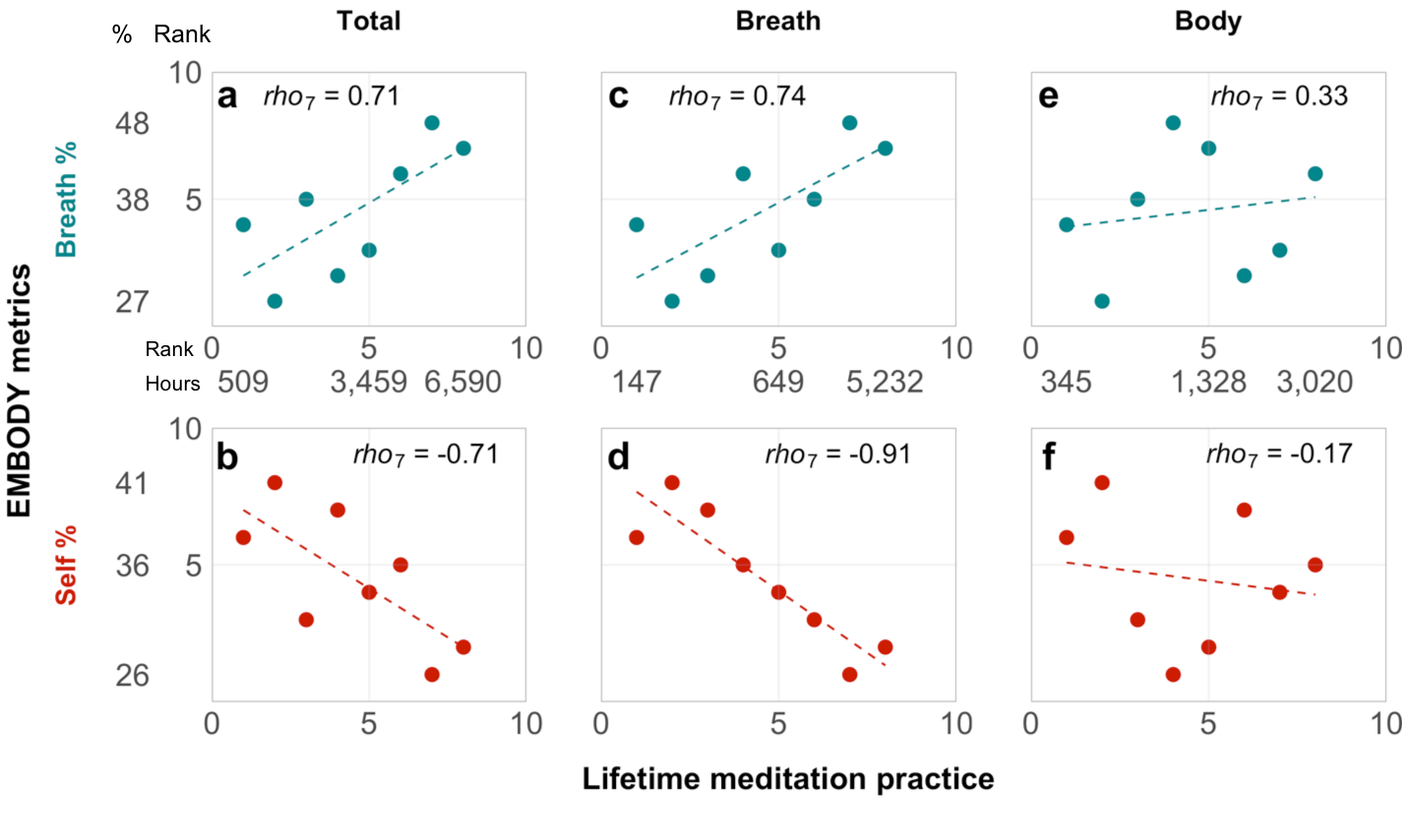

**Figure S3. Exploratory associations of mental states during meditation and lifetime meditation practice.** For exploratory purposes, meditation metrics from the EMBODY Task were non-parametrically correlated with lifetime hours of meditation practice in experienced meditators (N=8) using Spearman’s *rho* for descriptive purposes. Total lifetime hours of meditation practice in general were **(a)** positively associated with percentage time attending to the breath and **(b)** negatively associated with percentage time engaging in self-referential processing during meditation. Suggesting specificity, lifetime hours of meditating particularly on breath sensations were associated with **(c)** greater percentage time attending to the breath and **(d)** less percentage time engaged in self-referential processing during meditation, while hours meditating on other bodily sensations were less strongly associated with either **(e)** attending to breath or **(f)** self-referential processing. Ranks of both lifetime meditation hours and EMBODY metrics are displayed. Numerical values associated with ranks 1, 5, and 8 of each variable are displayed to aid interpretation of data.

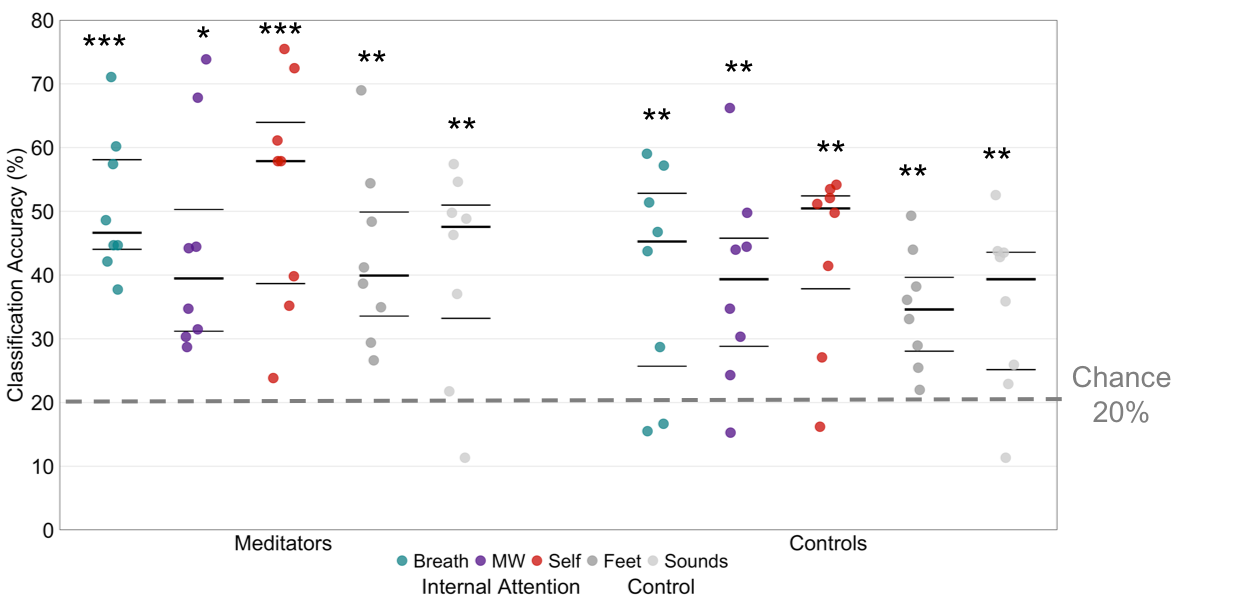

**Figure S4. EMBODY Step 1 classification accuracy by group.** Classifier accuracy of brain patterns associated with mental states from the Internal Attention task for each group. For both Meditators (N=8) and Controls (N=8), each brain pattern was recognized above theoretical chance levels (20%, indicated by dotted line). Beeswarm plots present each data point, the median (bold black line), and ±25^th^ percentile range (gray lines) of classifier accuracy.

* *p* < 0.05

** *p* < 0.01

*** *p* < 0.001

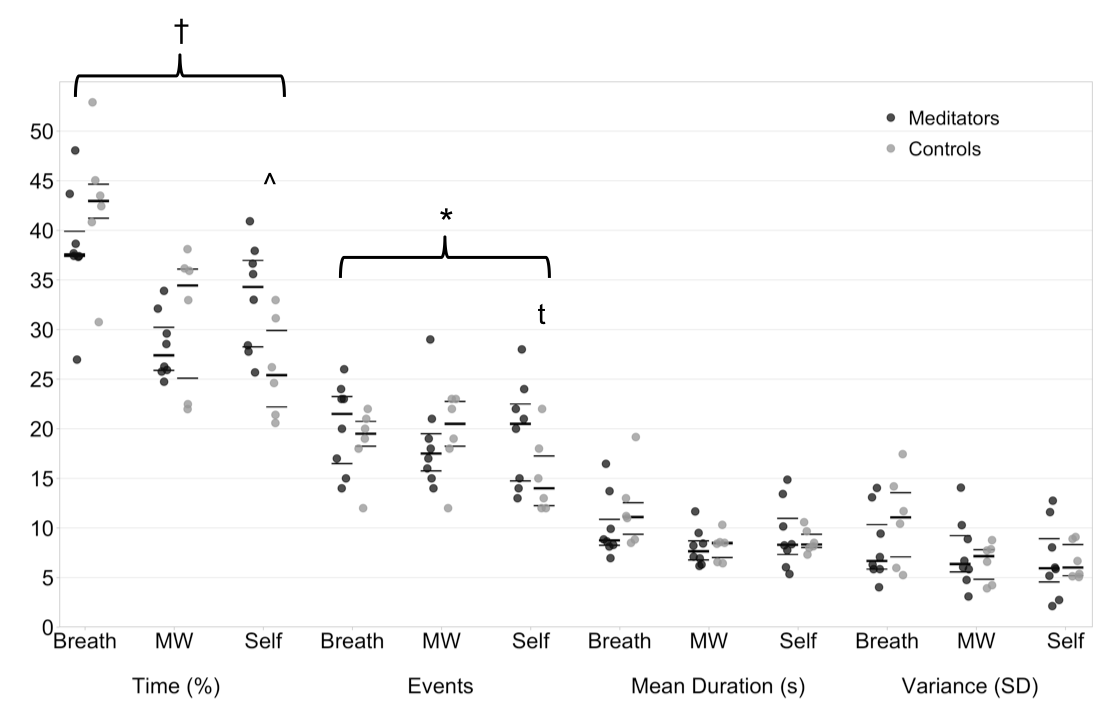

**Figure S5. Exploratory group differences in EMBODY meditation period metrics.** In initial exploratory analyses, we preliminarily compared Meditators (N=8) vs. Controls (N=6) in metrics from the 10-min period of breath meditation. Metrics were converted to ranks to account for small sample sizes, and the figure displays raw values for ease of interpretation. Beeswarm plots present each data point, the median (bold black line), and ±25^th^ percentile range (gray lines).

Significant and trend-level results are displayed to aid future research questions.

^†^ Group × Condition repeated measures ANOVA, *F_2_*_,12_ = 2.83, *p* = 0.08

^ Independent *t*_2,12_ = 2.66, *p* < 0.05

* Group × Condition ANOVA, *F*_2,12_ = 4.12, *p* < 0.05

^t^ Independent *t*_2,12_ = 1.74, *p* = 0.11

|  |  | **Group** |  |  |  |  |
| --- | --- | --- | --- | --- | --- | --- |
| **Demographics** |  | All | Final | Meditators | Controls | Final Controls |
| N |  | 16 | 14 | 8 | 8 | 6 |
| Age | Mean  (SD) | 38.31 (12.21) | 39.29  (12.55) | 38.38  (11.76) | 38.25  (13.48) | 40.5  (14.60) |
|  | Range | 25-63 | 27-63 | 28-61 | 25-63 | 27-63 |
| Gender | Female | 2 | 2 | 1 | 1 | 1 |
|  | Male | 12 | 10 | 6 | 6 | 4 |
|  | Non-binary | 2 | 2 | 1 | 1 | 1 |
| Race/Ethnicity^a^ | Asian/Asian-American | 1 | 1 | 0 | 1 | 1 |
|  | Hispanic/Latinx | 1 | 0 | 0 | 1 | 0 |
|  | White/Caucasian | 12 | 11 | 6 | 6 | 5 |
|  | Multiracial | 2^b^ | 2^b^ | 2^b^ | 0 | 0 |
| Income^c^ | Median level | 7  ($50,000-$59,999) | 7  ($50,000-59,999) | 6  ($40,000- $49,999) | 9  ($80,000-99,999) | 9  ($80,000- $99,999) |
|  | Range | 1-12  (<$10,000-$249,999) | 1-12  (<$10,000-$249,999) | 1-9  (<$10,000- $99,999) | 1-12  (<$10,000-$249,999) | 2-12  ($10,000-$249,999) |

**Table S1. Demographic information.** Demographics are reported for the full sample (N = 16), the final sample with distinguishable brain patterns (N = 14), all Meditators (N = 8), all Controls (N = 8), and final Controls with distinguishable brain patterns (N = 6).

^a^ Race and ethnicity were assessed as one category, so full race/ethnicity data are not available

^b^ 1 African American/White and 1 Asian/White

^c^ Participants were asked to report total household income over the past year by choosing 1 of 16 categories of income ranges. The categories are listed below:

1- Less than $10, 000
2- $10,000 - $14,999
3- $15,000 - $19,999
4- $20,000 - $29,999
5- $30,000 - $39,999
6- $40,000 - $49,999
7- $50,000 - $59,999
8- $60,000 - $79,999
9- $80,000 - $99,999
10- $100,000 - $149,999
11- $150,000 - $199,999
12- $200,000 - $249,999
13- $250,000 - $299,999
14- $300,000 - $399,999
15- $400,000 - $499,999
16- over $500,000

|  | **Meditators** | |  |
| --- | --- | --- | --- |
| **Meditation tradition(s)** | N | |  |
| Vipassana | 1 | |  |
| Zen | 2 | |  |
| Mixed | 1 Buddhist  1 Vipassana/Osho  1 Vipassana/Zen/Mahamudra/Dzogchen  1 Vipassana/Zen/Insight  1 Zen/Other | |  |
| **Lifetime Meditation Hours** | **Mean (SD)** | **Range** | **Years** |
| Total | 3495.1 (1930) | 509 - 6590 | 8 (3.66) |
| Breath | 1453.9 (1773) | 147 - 5232 |  |
| Body | 1410.1 (998) | 345 - 3020 |  |
| Other | 631.1 (536) | 0 - 1264 |  |

**Table S2. Meditation practice traditions and statistics.** Meditators reported the meditation traditions from which they received training. They reported lifetime meditation hours including total practice (Total), practice primarily focused on breath sensations (Breath), practice primarily focused on other body sensations (Body), and other meditation practices (Other, i.e., lovingkindness, mantra practice).

|  | **Breath** | **MW** | **Self** | **Feet** | **Sounds** | ***F*_4,79_** | ***p*** |
| --- | --- | --- | --- | --- | --- | --- | --- |
| **Roll** | 0.014 (.012) | 0.014 (.013) | 0.013 (.010) | 0.012 (.011) | 0.013 (.012) | 0.09 | 0.99 |
| **Pitch** | 0.031 (.014) | 0.034 (.020) | 0.030 (.012) | 0.027 (.011) | 0.028 (.012) | 0.61 | 0.66 |
| **Yaw** | 0.015 (.012) | 0.016 (.015) | 0.014 (.009) | 0.013 (.011) | 0.013 (.012) | 0.13 | 0.97 |
| **dS** | 0.048 (.035) | 0.050 (.034) | 0.043 (.025) | 0.040 (.024) | 0.040 (.029) | 0.40 | 0.81 |
| **dL** | 0.014 (.014) | 0.015 (.017) | 0.014 (.014) | 0.013 (.016) | 0.014 (.018) | 0.03 | 0.998 |
| **dP** | 0.061 (.022) | 0.055 (.019) | 0.054 (.016) | 0.049 (.014) | 0.049 (.014) | 1.35 | 0.26 |
| **Total** | 0.169 (.081) | 0.170 (.091) | 0.154 (.058) | 0.143 (.059) | 0.144 (.071) | 0.53 | 0.72 |

**Table S3. Head motion by Internal Attention task condition.** Mean (SD) head movement per volume in mm for each of 6 directions, as well as the total summed movement from all directions. Statistics reported for a one-way ANOVA between 5 conditions in each motion direction. Note that no differences in head motion between conditions were found.

dS = shifts in superior to inferior, dL = shifts in left to right, dP = shifts in posterior to anterior

|  | **Respiration Rate** | | | **Paired *t*_1,11_-tests** | | |
| --- | --- | --- | --- | --- | --- | --- |
|  | Breath | MW | Self | Br vs. MW | Br vs. Self | MW vs. Self |
| **Ppt 1** | 15.2 (1.8) | 16.7 (1.7) | 17.7 (1.2) | -3.65** | -5.50*** | -1.88 |
| **Ppt 2** | 11.9 (1.8) | 14.3 (1.4) | 13.9 (0.7) | -3.31** | -3.41** | 0.85 |
| **Ppt 3** | 18.8 (2.1) | 20.9 (1.2) | 20.1 (1.6) | -3.74** | -1.45 | 1.21 |

**Table S4.** Respiration rate in the IA task for 3 participants. Mean respiration rate (SD) for each condition relevant for meditation in the Internal Attention task (12 trials each), including paired *t*_1,11_-test values for each condition pair. * *p*<0.05, ** *p*<0.01, *** *p*<0.001

|  |  | **Decisions** |  |  |  |  |  |  |
| --- | --- | --- | --- | --- | --- | --- | --- | --- |
|  |  | Breath | MW | Self | Feet | Sounds | Accuracy | SD |
| **Condition** | Breath | **218.1** | 48.8 | 52.5 | *57.3* | 55.4 | 50.5**** | 16.6 |
|  | MW | 61.4 | **177.9** | *75.1* | 49.3 | 68.3 | 41.2*** | 18.2 |
|  | Self | 51.1 | 49.8 | **211.6** | *65.0* | 54.6 | 49.0**** | 18.4 |
|  | Feet | 55.8 | 51.1 | 65.8 | **186.5** | *72.8* | 43.2**** | 13.3 |
|  | Sounds | 60.5 | 59.6 | 54.3 | *69.0* | **188.6** | 43.7**** | 16.7 |

**Table S5. Internal Attention (IA) task classifier confusion matrix.** For all 16 participants in the IA task from Step 1, the mean decisions made in each category given each instructed condition are reported. **Bolded** numbers indicate the correct classifier decisions made for each instructed condition. *Italicized* numbers indicate the condition for which each instruction category was most likely to be “confused” with. Accuracy indicates the mean classifier accuracy for each instructed condition, and SD indicates the standard deviation of classifier accuracy.

******* *p* < 0.001 vs. 20% (theoretical chance)

******** *p* < 0.0001 vs. 20% (theoretical chance)

|  |  | **Internal Attention Neural Pattern** | | | | | | | | |
| --- | --- | --- | --- | --- | --- | --- | --- | --- | --- | --- |
|  |  | **All** (N = 16) | | | **Meditators** (N=8) | | | **Controls** (N=8) | | |
|  |  | Breath | MW | Self | Breath | MW | Self | Breath | MW | Self |
| *p* value | < 0.001 | 14 | 14 | 13 | 8 | 8 | 7 | 6 | 6 | 6 |
|  | < 0.01 | 15 | 14 | 13 | 8 | 8 | 7 | 7 | 6 | 6 |
|  | < 0.05 | 15 | 14 | 14 | 8 | 8 | 8 | 7 | 6 | 6 |
|  | > 0.05 | 1 | 2 | 2 | 0 | 0 | 0 | 1 | 2 | 2 |

**Table S6.** **Individual-level classification accuracies from the Internal Attention (IA) task.** The table indicates the number of participants for which each neural pattern relevant for breath-focused meditation (from Step 1 of the IA task) was recognized above chance within the individual. Individual-level accuracy was determined using a Chi-square test comparing the frequency of correct vs. incorrect MVPA decisions for each mental state compared to chance distribution (87 correct vs. 345 incorrect decisions). Participant frequencies are displayed for varying *p*-values. In order to use the neural patterns for subsequent decoding the meditation period (Step 2), we required that each individual have at least 2/3 brain patterns significantly identified above chance (*p* < 0.05). 14/16 participants survived this criterion, including all 8 Meditators and 6/8 Controls. Most of these individual-level brain patterns were distinguished at a high significance (*p* < 0.001).

See supplementary excel file “**TableS7.xlsx**”

**Table S7. Brain regions from group frequency importance maps.** Frequency maps were computed by summing each subject’s MNI-normalized importance map for breath, mind wandering (MW), and self-referential processing (Self) for All (**Fig. 3** from main manuscript), Positive, and Negative importance voxels (**Fig. S2**). Each voxel indicates the number of participants for which the voxel is important in distinguishing each mental state. “Positive importance” voxels had a positive weight and a positive *z*-scored average activation value (indicating that it was more active on average), and voxels with “negative importance” had a negative weight and a negative *z*-scored average activation value (indicating that it was less active on average; McDuff *et al.*, 2009). To identify important regions at the group level, maps were thresholded at $\geq$5 or 6 (at least half of the maximum frequency found in each mental state), and with a cluster size of 160 mm^3^ (at least 20 contiguous voxels). The MNI coordinates and value of the peak frequency voxel are reported. If peak voxels consisted of multiple contiguous voxels, the coordinates for the central voxel in each sub-cluster were reported. PFC = prefrontal cortex.

^A^ = All importance cluster region

^N^ = Negative importance cluster region

^P^ = Positive importance cluster region

|  | **EMBODY Meditation Period Metrics – Mean (SD)** | | | | | |
| --- | --- | --- | --- | --- | --- | --- |
|  | Percentage Time | Number of events | Mean duration | Variance (SD) | Total events | Distraction from breath |
| **All (N=14)** |  |  |  |  | 56.4 (11.3) | 20.4 (4.8) |
| Breath | 40.2 (6.6) | 19.6 (4.0) | 10.9 (3.5) | 9.3 (4.2) |  |  |
| MW | 29.6 (5.3) | 19.0 (4.4) | 8.1 (1.6) | 7.1 (2.9) |  |  |
| Self | 30.2 (6.3) | 17.8 (5.1) | 9.0 (2.6) | 6.8 (3.0) |  |  |
| **Meditators (N=8)** |  |  |  |  | 58.5 (13.6) | 19.8 (4.6) |
| Breath | 38.4 (6.1) | 20.3 (4.5) | 10.1 (3.3) | 8.2 (3.6) |  |  |
| MW | 28.3 (3.3) | 18.6 (4.7) | 8.0 (1.9) | 7.5 (3.5) |  |  |
| Self | 33.2 (5.5) | 19.6 (5.3) | 9.3 (3.4) | 6.8 (3.8) |  |  |
| **Controls (N=6)** |  |  |  |  | 53.5 (7.4) | 21.1 (5.3) |
| Breath | 42.6 (7.2) | 18.7 (3.6) | 12.0 (3.9) | 10.8 (4.7) |  |  |
| MW | 31.3 (7.2) | 19.5 (4.2) | 8.1 (1.4) | 6.5 (2.0) |  |  |
| Self | 26.2 (5.0) | 15.3 (4.0) | 8.7 (1.2) | 6.7 (1.9) |  |  |

**Table S8. EMBODY Task meditation period metrics.** Means and SDs for meditation period metrics for the All participants with distinguishable brain patterns used to decode the meditation period (N = 14/16), Meditators (N = 8), and Novice Controls (N = 6). Mean duration, variance of mean duration (SD), and distraction from breath are reported in seconds.

|  | **Analysis** | | | | **Means** | | | **Paired *t*_1,13_-tests** | | |
| --- | --- | --- | --- | --- | --- | --- | --- | --- | --- | --- |
|  | Smoothed | Min event length (s) | | % Data excluded | Breath | MW | Self | Br-MW | Br-Self | MW-Self |
| **Percentage Time** | 0 | | 0 | 0.0 | 38.44 | 30.87 | 30.69 | 3.55** | 3.17** | 0.09 |
|  | 0 | | 2 | 8.9 | 39.39 | 30.25 | 30.36 | 3.67** | 3.26** | -0.05 |
|  | 0 | | 3 | 17.0 | 40.16 | 29.62 | 30.22 | 3.84** | 3.22** | -0.23 |
|  | 0 | | 4 | 23.6 | 40.65 | 28.86 | 30.49 | 4.12** | 3.06** | -0.60 |
|  | 1 | | 2 | 6.3 | 39.47 | 30.28 | 30.25 | 3.73** | 3.29** | 0.02 |
|  | **1** | | **3** | **15.7** | **40.22** | **29.61** | **30.17** | **3.87**** | **3.18**** | **-0.24** |
|  | 1 | | 4 | 22.7 | 40.68 | 28.89 | 30.43 | 4.15** | 3.04** | -0.58 |
| **Number of Events** | 0 | | 0 | 0.0 | 46.07 | 49.21 | 46.00 | -1.57 | 0.04 | 1.91 |
|  | 0 | | 2 | 8.9 | 23.43 | 23.93 | 23.07 | -0.49 | 0.33 | 0.82 |
|  | 0 | | 3 | 17.0 | 16.21 | 16.57 | 15.43 | -0.48 | 0.84 | 1.22 |
|  | 0 | | 4 | 23.6 | 12.86 | 12.21 | 11.86 | 0.73 | 1.61 | 0.38 |
|  | 1 | | 2 | 6.3 | 23.43 | 23.93 | 23.07 | -0.49 | 0.33 | 0.82 |
|  | **1** | | **3** | **15.7** | **16.21** | **16.57** | **15.43** | **-0.48** | **0.84** | **1.22** |
|  | 1 | | 4 | 22.7 | 12.86 | 12.21 | 11.86 | 0.73 | 1.61 | 0.38 |
| **Mean Duration** | 0 | | 0 | 0.0 | 5.33 | 3.91 | 4.11 | 3.64** | 2.74* | -0.80 |
|  | 0 | | 2 | 8.9 | 9.67 | 7.07 | 7.52 | 3.55** | 2.22* | -0.87 |
|  | 0 | | 3 | 17.0 | 13.12 | 9.25 | 10.54 | 3.72** | 2.37* | -1.72 |
|  | 0 | | 4 | 23.6 | 15.53 | 11.56 | 13.19 | 2.95* | 1.69 | -1.27 |
|  | 1 | | 2 | 6.3 | 9.95 | 7.28 | 7.71 | 3.62** | 2.25* | -0.81 |
|  | **1** | | **3** | **15.7** | **13.33** | **9.39** | **10.70** | **3.75**** | **2.35*** | **-1.69** |
|  | 1 | | 4 | 22.7 | 15.70 | 11.72 | 13.33 | 2.92* | 1.65 | -1.21 |
| **SD** | 0 | | 0 | 0.0 | 6.52 | 4.62 | 4.90 | 2.59* | 2.40* | -0.74 |
|  | 0 | | 2 | 8.9 | 9.66 | 6.69 | 6.64 | 2.49* | 2.48* | 0.09 |
|  | 0 | | 3 | 17.0 | 11.70 | 8.30 | 7.91 | 2.34* | 2.96* | 0.43 |
|  | 0 | | 4 | 23.6 | 12.72 | 9.44 | 9.26 | 1.85 | 2.83* | 0.11 |
|  | 1 | | 2 | 6.3 | 10.02 | 6.93 | 6.85 | 2.52* | 2.49* | 0.15 |
|  | **1** | | **3** | **15.7** | **12.01** | **8.51** | **8.08** | **2.36*** | **2.97*** | **0.45** |
|  | 1 | | 4 | 22.7 | 12.99 | 9.71 | 9.43 | 1.79 | 2.72* | 0.16 |

**Table S9. Meditation metrics from alternate analyses of the meditation period.** Meditation period decisions analyzed with varying “mental event” lengths (2, 3, or 4s), with or without a smoothing algorithm, which re-labels a single incongruous decision between two events of the same type (e.g., MW event – Self decision – MW event) according to the category of the surrounding events (e.g., Self => MW; 0 indicates no smoothing algorithm used, 1 indicates that a single incongruous decision was smoothed). Data were smoothed according to the corresponding event length type (e.g., if the analysis had a minimum event length of 2s, data were smoothed between events that were 2s long). To test the stability of the main finding where percentage time engaged in breath attention was greater than mind wandering or self, we re-computed planned paired t-tests between Breath vs. MW, Breath vs. Self, and MW vs. Self for completeness. Note that for percentage time of mental states during meditation, every analysis yields the main finding that participants spend more time engaged in Breath vs. MW or Self. **Bold** indicates the analysis presented in the main manuscript (smoothing present, minimum event length=3s).

* *p*<0.05, ** *p*<0.01

|  |  | **EMBODY metrics** | |  |
| --- | --- | --- | --- | --- |
| **Survey** | **Sub-scale** | **Breath %** | **MW %** | **Self %** |
| *A priori* |  |  |  |  |
| MAIA | Noticing | -0.32 | -0.06 | 0.42 |
|  | Attention Regulation | -0.58 | 0.10 | 0.52 |
| FFMQ^a^ | Observing | -0.39 | 0.54 | 0.15 |
|  | Acting with Awareness | -0.45 | 0.30 | 0.35 |
| *Exploratory* |  |  |  |  |
| MAIA | Not Distracting | -0.63 | -0.02 | 0.73 |
|  | Not Worrying | -0.29 | 0.07 | 0.44 |
|  | Emotional Awareness | -0.34 | 0.12 | 0.32 |
|  | Self-Regulation | -0.38 | 0.05 | 0.37 |
|  | Body Listening | -0.42 | -0.05 | 0.54 |
|  | Trusting | -0.36 | 0.01 | 0.42 |
| FFMQ^a^ | Describing | -0.41 | 0.28 | 0.34 |
|  | Nonjudging | -0.40 | 0.04 | 0.46 |
|  | Nonreacting | -0.31 | -0.09 | 0.56 |

**Table S10.** **Exploratory associations between meditation metrics and trait interoception and mindfulness.** Percentage time engaged in mental states during meditation from the EMBODY Task was correlated descriptively with trait questionnaires of interoception and mindfulness (respectively, MAIA: Multidimensional Interoceptive Awareness Inventory(Mehling *et al.*, 2012) and FFMQ: Five Facet Mindfulness Questionnaire (Baer *et al.*, 2006)). Strongest *a priori* hypotheses for associations were with the MAIA Noticing and Attention Regulation scales and FFMQ Observing and Acting with Awareness scales (see **SI-Methods**). Correlations are reported for exploratory purposes only.

^a^ Full FFMQ data were collected for 13/16 participants, ranging from 2-18 months (mean=12) after the brain scan. 11 participants had valid EMBODY metrics and full FFMQ scores that could be used for correlation tests. The original FFMQ was mistakenly administered with only 14/39 total items and was not used for analyses. These data should be interpreted with caution.
